# Extending species-area relationships into the realm of ecoacoustics: The soundscape-area relationship

**DOI:** 10.1101/2023.02.08.527658

**Authors:** Thomas Luypaert, Anderson S. Bueno, Torbjørn Haugaasen, Carlos A. Peres

## Abstract

The rise in species richness with area is one of the best-studied patterns in ecology. Yet, little is known about the spatial scaling of alternate dimensions of diversity. Here, we provide empirical evidence for a relationship between the richness of acoustic traits emanating from the landscape, or soundscape richness, and the island area, which we term the Soundscape Area Relationship (SSAR). We show a positive relationship between the gamma soundscape richness and island area, with slope and R^2^-values of comparable magnitude to those reported in the literature. This relationship breaks down at the smallest spatial scales, indicating a small-island effect. Moreover, we demonstrate a positive spatial scaling of the plot-scale alpha soundscape richness, but not the beta soundscape turnover, suggesting disproportionate effects are an important underlying mechanism. We conclude that the scaling of biodiversity can be extended to the realm of ecoacoustics, implying soundscape metrics are sensitive to fundamental ecological patterns and useful to disentangle the complex mechanisms that drive them.

## 1. Introduction

Humankind’s pursuit of improved welfare, food security, and protection against the natural world has led to the rapid and widespread destruction, degradation and fragmentation of natural ecosystems (Haddad et al. 2015; Newbold et al. 2015). Driven by an interest in the ecological mechanisms underlying species assembly in insular landscapes and, more recently, by increasing concerns about the effects of habitat fragmentation, biodiversity research in patch-matrix landscapes has been a fundamental area of research for over a century (Giladi et al. 2014; Jacquet et al. 2017). This topic is often addressed through the theory of island biogeography (IBT; MacArthur and Wilson 1967), where habitat patches are perceived as islands in a hostile non-habitat matrix. Per the IBT, the biological processes of species richness regulation in insular systems are modulated by the interplay between patch size and isolation. These factors influence metacommunity dynamics such as migration and extinction in habitat patches, leading to two foundational observations: (i) smaller habitat patches have a reduction in the patch-scale species richness (species-area relationship or SAR); and (ii) more isolated habitat patches have a reduced patch-scale species richness (species-isolation relationship or SIR; Giladi et al. 2014).

The spatial scaling of species richness with area (SAR) is one of the oldest, best-documented, and most ubiquitous ecological patterns (Arrhenius 1921; Gleason 1922; Connor and McCoy 1979; Rosenzweig 1995; Lomolino 2000; Drakare et al. 2006; Matthews et al. 2016). Although SARs can take many different forms, here we will focus on the Island Species-Area Relationship (ISAR or Type IV SAR), which quantifies the increase in species richness with increasing island or habitat patch size (Scheiner 2003). The relationship is typically positive, with the exception of particularly small islands, where stochastic processes may mask the effect of area on the species richness (the small island effect; Niering 1963; Woodroffe 1986; Lomolino 2000). Yet, despite its importance to conservation biogeography, there remains a great deal of ambiguity regarding the underlying mechanisms that shape, regulate and maintain SARs across space (Scheiner et al. 2011; Chase et al. 2019; Gooriah 2020).

The null hypothesis is that ISARs are the result of sampling effects. Firstly, it is possible that ISAR patterns are produced by sampling artefacts (Preston 1962; Schoereder et al. 2004), whereby larger islands need a higher sampling effort to describe the species richness adequately. In doing so, more individuals are sampled, which, by probability, increases the likelihood of more species being sampled (Hill et al. 1994). Secondly, passive sampling effects dictate that the species richness in island systems is controlled by ecological sampling processes, by which larger areas represent a larger sample of the original species pool than smaller ones, and by probability, sample more species (Chase et al. 2019). Although sampling artefacts and passive sampling capture similar processes, the first represents a human-introduced data error generated by the sampling procedure, whereas the second reflects real differences in species richness between islands resulting from ecological sampling processes.

Alternatively, it has been hypothesised that ISARs are the result of the biological differences linked to island size. According to the theory of disproportionate effects, larger islands contain more species because patch size affects the biological processes of species richness regulation (Schoereder et al. 2004). According to the IBT, extinction rates are inversely proportional to the population size, and the number of individuals a patch can sustain is proportional to the patch size. Hence, a reduced patch size leads to increased extinction rates and reduced patch-scale species richness. Moreover, smaller patches are believed to have reduced colonisation rates (through target effects), leading to higher extinction rates and lower species richness. Moreover, disproportionate effects can emerge through a range of other mechanisms, including edge-effects and a reduction in trophic levels (see Chase et al. 2019 for a full overview). Finally, the theory of heterogeneity effects posits that the compositional heterogeneity of species on islands is influenced by island size. For instance, if larger islands contain a larger variety of unique habitats, with each habitat containing a distinct set of associated species, more habitats and accompanying species are sampled for large islands, and thus the number of species increases with area (Williams 1964).

In contrast to SARs, evidence for the species-isolation relationship (SIR) is more scarce (but see Giladi et al. 2014). According to the IBT, the SIR can be attributed to the effects of decreasing immigation, and thus, colonisation rates from the *‘mainland’* to isolated patches, leading to reduced rescue effects, higher rates of extinction, and ultimately, a lower species richness (Helmus et al. 2014). In patchy habitat systems, however, immigrants travel predominantly from other nearby habitat patches rather than a mainland area (Fahrig 2013). Hence, the isolation of a patch depends on the amount of nearby habitat within a radius around the patch, and thus, the isolation can be seen as a measure of local landscape-scale habitat amount. This concept is built upon by Fahrig (2013), who proposes that, in fragmented landscapes, habitat patches do not behave as closed entities, but allow travel between patches across the matrix to a varying degree. Consequently, the author states that the IBT’s observed patch-scale SARs and SIRs are just reflections of the sample area effect: larger and less isolated patches tend to be surrounded by a larger local landscape-scale habitat amount, which contains more individuals, and by extension, more species (the habitat amount hypothesis or HAH; Fahrig et al. 2013).

In summary, the importance of the patch-scale habitat amount and isolation (following the IBT) versus the local landscape-scale habitat amount (following the HAH) will depend on the degree to which patches behave as closed units to the inhabiting community, which is determined by the hostility of the matrix and the dispersal ability of the taxonomic group under investigation. In reality, it is likely that multiple mechanisms are contributing to the observed spatial biodiversity patterns that are reported in the literature. However, to date, elucidating which ecological mechanisms are at work has proven challenging. Additionally, investigations into spatial biodiversity patterns in insular systems have traditionally been limited to quantifications of taxonomic species richness. Yet, biological diversity clearly transcends species richness alone. Still, the spatial variation of alternate dimensions of biodiversity remains a gap in the literature (Delsol et al. 2018; Galiana et al. 2022; de Camargo et al. 2019; Gonzalez et al. 2020).

One of these alternative aspects of diversity can be found in the field of ecoacoustics, which makes use of the soundscape, or the combination of all sounds produced in an environment, to make inferences about ecological processes at a landscape scale (Farina & Gage 2017). The premise is that, when ecological processes or disturbances lead to changes in an ecosystem and its inhabitants, this change will be reflected by the sounds emanating from the landscape (Stowell & Sueur 2020). However, rather than deriving taxonomic information by isolating and identifying species’ calls from sound files, which can be prohibitively time-consuming for large acoustic datasets, eco-acoustic methods detect ecological signals by making use of acoustic indices (Sueur et al. 2014). These mathematical formulae extract information on the diversity of acoustic traits emanating from the landscape by quantifying the amplitude variation across the time-frequency domain of sound files (Eldridge et al. 2018). If these metrics respond in a consistent and predictable manner to underlying changes in the ecosystem, acoustic indices can be used to lift the veil on landscape-scale processes (Bradfer-Lawrence et al. 2020). Indeed, acoustic indices have been applied successfully to answer questions in biogeography such as identifying habitat disturbances (e.g. Burivalova et al. 2018), distinguishing habitat types (e.g. Bormpoudakis et al. 2013), assessing landscape configuration (e.g. Fuller et al. 2015), and more. Yet, other than a pioneer study by Müller et al. (2020), surprisingly, the spatial scaling of eco-acoustic metrics and the mechanisms that govern them in space have received little attention.

Thus, in this study, we examined the spatial variation in soundscape richness using an insularized tropical rainforest system. In doing so, we aimed to answer the following research questions:

1. What is the relative importance of the patch-scale habitat amount (or island size - SAR) versus the isolation (or the inverse the landscape-scale habitat amount - SIR) on the soundscape richness?
2. If a relationship between the soundscape richness and island size is observed, what are the mechanisms driving this relationship?
  a. Is there a relationship between the island-wide gamma soundscape richness and island size?
    i. If so, can we observe a small-island effect?
    ii. How do the slope and and R^2^-values compare to other studies in the area?
  b. Does the plot-scale alpha soundscape richness scale with island size?
  c. Does the beta soundscape turnover scale with island size?

To answer these questions, we collected long-duration acoustic recordings for a set of Amazonian land-bridge islands at the Balbina Hydroelectric Reservoir in Central Amazonia. We sampled 72 plots on 51 islands ranging from 0.45 – 668 ha. For each island, we calculated the island-wide soundscape richness (gamma) using the metric described in Luypaert et al. (2022) and assessed its relationship with island size and isolation. Furthermore, to disentangle the mechanisms underlying potential SARs, we employed an adapted version of the multi-scale framework proposed by Chase et al. (2019), decomposing the soundscape richness into its alpha and beta components and evaluating their relationship with island size.

Together, these questions will yield insights into the factors that govern acoustic diversity in fragmented landscapes and shed light on the use of eco-acoustic metrics as a tool to disentangle complex ecological mechanisms.

## 2. Methods

### 2.1. Data collection

#### 2.1.1. A multi-scale sampling protocol to uncover spatial biodiversity patterns

Much of the contention relating to the ecological mechanisms driving biodiversity patterns in insular systems is linked to methodological inconsistencies in how these patterns should be quantified (Chase et al. 2019; Gooriah 2020). For instance, historically, most studies have investigated ISARs using uniform sampling protocols (a fixed sampling effort across the island area gradient), characterizing the relationship between the island-wide gamma richness and island size (Schoereder et al. 2004; Chase et al. 2019). Although this approach is useful in capturing the shape of the ISAR while dealing with confounding non-biological sampling processes that influence ISAR patterns (sampling artefacts), it can tell us little about the potential ecological mechanisms driving the observed patterns. Equalizing the sampling effort for varying island sizes may obscure variations in habitat diversity, and thus miss out on important beta diversity effects (e.g. heterogeneity effects; Schoereder et al. 2004). Moreover, by focussing on island-wide patterns in biological richness, local-scale (alpha) biodiversity patterns are overlooked (*e*.*g*. disproportionate effects).

To elucidate which ecological mechanisms underlie the spatial biodiversity patterns in our study system, we employed a modified version of the generalized multi-scale and multi-metric framework outlined in Chase et al. (2019). We adopted a spatial sampling design consisting of multiple standardized acoustic sampling plots per island, with the number of plots per island being directly proportional to the island size (Fig. 1C). By pooling the sampled plots per island, we can derive the island-wide gamma soundscape richness. This metric can be used to assess the relative importance of the patch-scale habitat amount (island size) versus the landscape-scale habitat amount (isolation).

**Figure 1:**
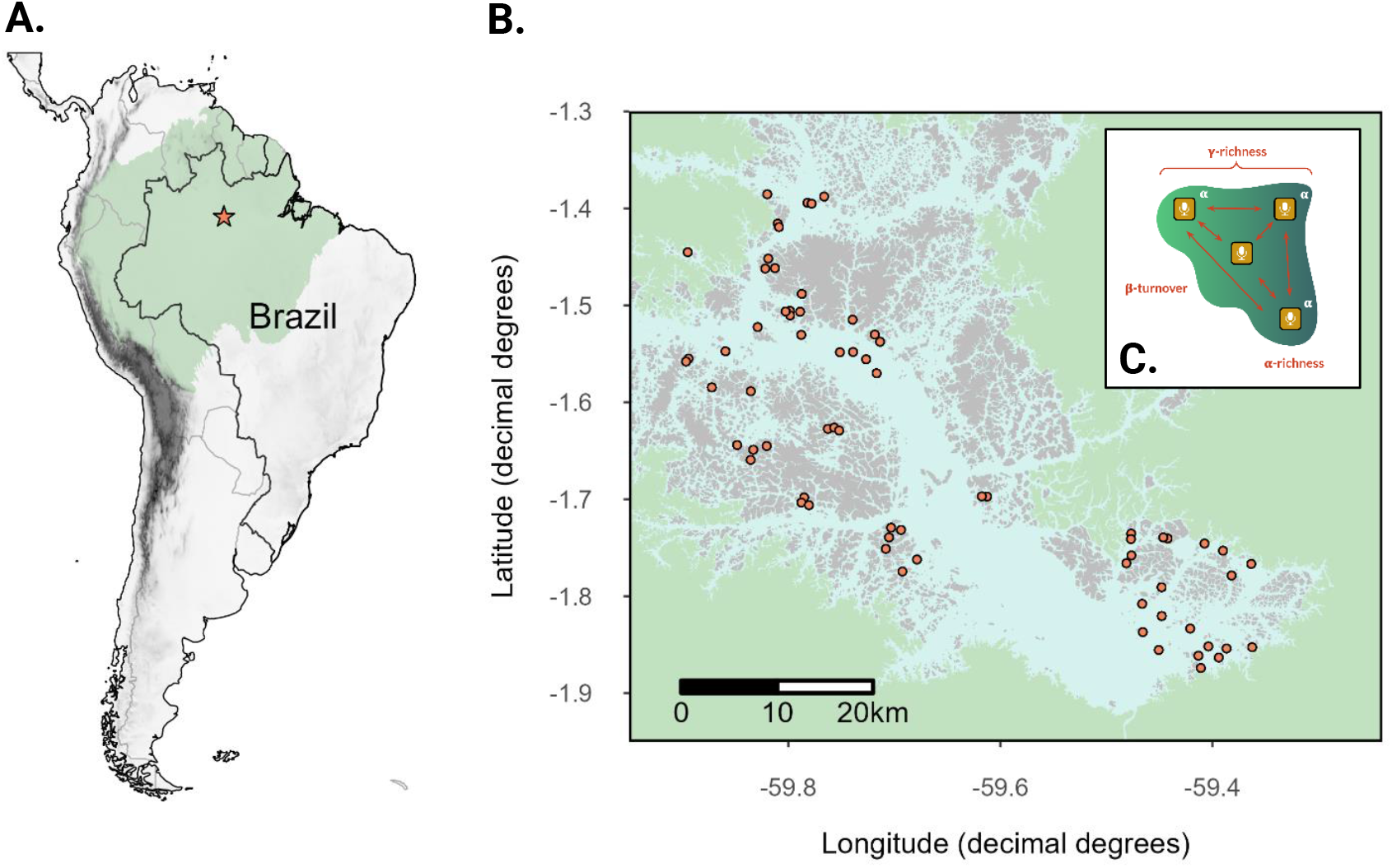
**A**. Location of the Balbina Hydroelectric Reservoir (BHR; orange star) in the central Amazonian region (light green) of Brazil. **B**. A detailed overview of the BHR (blue) showing over 3500 hilltop islands (grey), surrounded by continuous forest (green). For this study, 69 sampling sites (orange) on 49 islands were sampled. **C**. An overview of the spatial sampling design employed in this study. The green area represents an island, with multiple acoustic sampling plots in yellow. This sub-island scale sampling design allows to quantify not only the island-wide gamma soundscape richness, but also the local-scale alpha soundscape richness and beta turnover components.

Moreover, we can look at the soundscape richness at sub-island scale to shed light on the ecological mechanisms governing ISARs in our study area. To account for the potentially confounding effects of sampling artefacts, we can compare both the unrarefied and rarefied gamma soundscape richness. Moreover, to discern whether the soundscape-area relationship is not just the result of passive sampling, we can investigate the soundscape richness at local plot scales (alpha soundscape richness) and between-plot turnover (beta soundscape turnover). If remnant size affects the processes of biodiversity regulation (disproportionate effects), such as migration-extinction dynamics, we would expect the local-scale alpha soundscape richness to correlate positively with the island size. Finally, if ISARs are the result of higher habitat diversity on larger islands (heterogeneity effects), we would expect the between-plot soundscape turnover to correlate positively with island size.

#### 2.1.2. Acoustic data collection

Acoustic data were collected at the Balbina Hydroelectric Reservoir (BHR) in Brazilian Amazonia (Fig. 1A-B). The BHR is one of the largest hydroelectric reservoirs on Earth and was formed when the Uatumã River, a tributary of the Amazon, was dammed in 1987 (Benchimol and Peres 2015a). This flooding event turned the hilltops of the former primary continuous forest into > 3,500 islands covering an area of approximately 300,000 ha, and ranging in size from 0.2 to 4,878 ha (Bueno and Peres 2019).

Long-duration acoustic surveys were conducted at the BHR between July and December 2015, sampling 74 forest islands (see Bueno et al. 2020). We adopted a proportional sampling scheme, for which the number of sampling plots per island increased with island size and ranged from 1 to 7 plots per island (see S1 for details). At each sampling plot, a passive acoustic sensor was deployed on a tree trunk at 1.5 m height from the ground with the microphone pointing downward. Each acoustic sensor consisted of an LG smartphone enclosed within a waterproof case with an external connector linked to an omnidirectional microphone and was set to record 1 min / 5min at a sampling rate of 44.1 kHz for 4-10 days using the ARBIMON Touch application (ARBIMON, https://arbimon.rfcx.org/). Due to poor recording quality, and to retain a proportional sampling scheme, several sampling sites were removed from the study (see S1). Ultimately, 69 sampling plots (1-4 plots per island) on 49 islands (0.45 - 668.03 ha) were retained in the study (Fig. S1; Table S1).

### 2.2. Calculating model variables

#### 2.2.1. Response variable - a metric of soundscape richness

To quantify the diversity of acoustic traits emanating from the landscape, we followed the analytical pipeline outlined in Luypaert et al. (2022) to calculate the *soundscape richness* acoustic index. This metric has previously been shown to attain a strong positive correlation with the species richness of sound-producing organisms in the BHR (Luypaert et al. 2022). As eco-acoustic metrics aim to capture ecological patterns without the need to isolate and identify species’ vocalizations from sound files, they generally lack a unified unit of diversity measurement (e.g. species). To overcome this, the pipeline quantifies the richness of Operational Sound Units (OSUs), a unit of measurement that groups sounds by their shared acoustic traits in the time-frequency domain of the acoustic space in which species produce sound.

Briefly, first, we equalized the sampling effort across all sampling plots to 5 sampling days. Next, we calculated the acoustic cover (CVR) spectral index for each 1-min sound file at every plot using a sampling rate of 44,100 Hz and a window length of 256 frames. For each plot, the spectral CVR-index files were concatenated chronologically into a time-by-frequency data frame of CVR-index values. Then, we determined the detection/non-detection of Operational Sound Units (OSUs) per 24h sample of the soundscape by converting the raw CVR-index values into a binary variable using the *‘IsoData’* binarization algorithm. In doing so, per plot, we obtained an OSU-by-sample incidence matrix, capturing the detection/non-detection of OSUs for each 24h soundscape sample in the 5-day acoustic survey. This matrix forms the base of all subsequent soundscape richness computations.

To quantify the overall island-wide gamma soundscape richness, per island, we pooled the OSU-by-sample incidence matrices across all sampling plots on that island. Next, we calculated the unrarefied island-wide gamma soundscape richness by counting the number of unique OSUs that were detected on each island. This metric forms the base of all modelling procedures in section 2.5. Moreover, in section 2.6., we investigated the spatial scaling of soundscape richness at sub-island scales using the: (i) rarefied gamma soundscape richness; (ii) alpha soundscape richness; and (iii) beta soundscape turnover.

#### 2.2.2. Predictor variables

The metric of island size used here corresponds to the total forest cover per island. We computed the metric using QGIS (QGIS Association 2022), employing a pre-classified image obtained from Landsat (30m resolution; collection 2, 2015, Amazon; available at http://mapbiomas.org). Forest cover was calculated as the amount of ‘*dense forest’* per island (pixel value 3), as other pixel values contained either heavily degraded or non-forest cover types. Using this classification type, the area of forest habitat was derived per island. For more information, consult Bueno et al. (2020).

For the isolation metric, we followed MacDonald et al. (2018), using QGIS to calculate island isolation as the proportion of water (1 – proportion of land area) within a range of buffer sizes calculated from the island edge. To determine the optimal scale-of-effect for our isolation variable (see Jackson and Fahrig 2015), we trialled 40 different buffer sizes (50-2000m at 50m intervals), choosing the spatial scale at which the isolation metric attains the strongest relationship with soundscape richness. The highest correlation between the soundscape richness and our isolation metric was attained at a scale-of-effect of 650 m (see S3 for detailed overview).

### 2.3. The effect of patch-scale versus landscape-scape habitat amount on soundscape richness (RQ1)

Despite the debate surrounding the most suitable modelling procedure (see Tjørve 2003, 2012; Triantis et al. 2012), ISARs are most often mathematically approximated by the power law function (Arrhenius 1921), defined as *S = cAz*, where *S* is the number of species/biodiversity units, *A* is the area of the habitat patch, and *c* and *z* represent the slope and intercept of the equation in log-log space. The power-law model has previously been shown to provide the best fit for ISARs at intermediate spatial scales such as the one in our study system (He and Legendre 1996; Triantis et al. 2012; Matthews et al. 2016). Moreover, later on, we aim to compare the strength and slope of a potential soundscape-area relationship to conventional ISAR studies for a range of taxonomic groups in the study area, which were mostly fitted using the power-law model. As such, for comparative purposes, we chose to fit all models including island-scale habitat amount using a power-law model framework. All power-law models were fitted using the ‘*lin_pow’* function of the ‘*sars*’ R-package (v1.3.5 - Matthews et al. 2019) using a log10 transformation.

In addition to the patch-scale habitat amount (or island size), we were also interested in the influence of island isolation (measured as the inverse of the landscape-scale habitat amount). To assess the relative importance of these two predictor variables, first, we used partial regression plots to visually explore each variable’s influence on the unrarefied soundscape richness while accounting for the variation taken up by the other variable. Next, we assessed whether an interaction effect was present between both predictor variables using conditioning scatterplots. Finally, we fitted a series of linear models and used an information theoretic approach for model selection (Burnham and Anderson 2004):

1. *log*_*10*_*(gamma soundscape richness) ~ log*_*10*_*(island area) + isolation + log*_*10*_*(island area)*isolation*
2. *log*_*10*_*(gamma soundscape richness) ~ log*_*10*_*(island area)*
3. *log*_*10*_*(gamma soundscape richness) ~ log*_*10*_*(island area) + isolation*
4. *log*_*10*_*(gamma soundscape richness) ~ isolation*
5. *log*_*10*_*(gamma soundscape richness) ~ 1*

As we know our predictor variables are correlated with each other (see S4.2), we checked for multicollinearity between the predictors by calculating the Variance Inflation Factor (VIF) using the ‘*vif’* function from the ‘*car’* R-package (Fox and Weisberg 2019 - version 3.1-0). We observed a VIF of 1.44, which is within the acceptable range to retain both predictor variables in the model (Johnston et al. 2018). For each model, we tested whether the following assumptions were met: (i) a normal distribution of residuals; (ii) homoscedasticity of residuals; (iii) a zero-mean of residuals; and (iv) independence of residual terms. For a detailed overview of the modelling procedure, consult supplementary material S4.

### 2.4. Decomposing the ecological mechanisms underlying ISARs (RQ2)

#### 2.4.1. Testing for a small-island effect

Due to the influence of non-area-related stochastic effects, species-area relationships are known to break down at very small scales, a phenomenon known as the *small island effect*. As such, prior to proceeding with subsequent analyses, we first checked for the presence of a small island effect by comparing four ISAR model types for the unrarefied gamma soundscape richness using the ‘*sar_threshold’* function from the ‘*sars’* R-package (v1.3.5 - Matthews et al. 2019): (i) a continuous one-threshold model; (ii) a left-horizontal one-threshold model; (iii) a power-law model without a small island effect; and (iv) an intercept-only model. We performed model selection using several model selection metrics (small-sample correct Akaike Information Criterion (AICc), Bayesian Information Criterion (BIC) and adjusted R^2^). If a small island effect was detected, we used the threshold value as a cut-off, excluding all island below the threshold value from subsequent analyses investigating soundscape richness-area patterns (see supplementary material S5 for detailed breakdown).

#### 2.4.2. The effect of sampling artefacts on the soundscape richness ~ island size relationship

As previously indicated, biological richness may increase with island area simply because of sampling artefacts (Schoereder et al. 2004). In the context of our workflow, the proportional sampling scheme we adopted may lead to more OSUs being detected because larger islands were sampled more intensely. To account for this unequal sampling effort between islands, we adopted a rarefaction procedure to calculate the rarefied gamma soundscape richness per island. Here, we employed both temporal-effort-based and plot-based rarefaction procedures to calculate the rarefied gamma soundscape richness, rarefying the sampling effort of the pooled OSU-by-sample incidence matrix to five sampling days and 1 plot per island respectively (see supplementary material S2 for detailed outline). Using the power-law model, the relationship between the rarefied gamma soundscape richness and island area was quantified.

#### 2.4.3. Alpha soundscape richness (α)

To test the role of disproportionate effects in generating the SSAR, we assessed the relationship between the plot-scale alpha soundscape richness and island area. We sub-sampled the total dataset to obtain a uniform sampling regime consisting of one plot per island, ensuring all plots had equal weight on the final alpha soundscape richness-area relationship by repeating the sub-sampling process until all possible combinations of 1-plot subsets across the islands in the study were generated (# subsets = 110,592). Then, we quantified the relationship between the alpha soundscape richness and island area by fitting a power-law model to the total of all one-plot-per-island subsets.

#### 2.4.4. Beta soundscape turnover (β)

To assess whether heterogeneity effects were driving the SSAR, we assessed the relationship between the beta soundscape turnover and island area. We calculated the beta soundscape turnover per island using the multiplicative framework offered by Hill numbers, whereby beta is calculated by dividing the regional (unrarefied island-wide gamma) soundscape richness by the average local (alpha) soundscape richness. The beta turnover captures the degree of heterogeneity in the OSU composition across plots and ranges from 1 to N, with N being the number of plots per island. The beta turnover can be seen as the effective number of completely distinct soundscapes per island (Tuomisto 2010).

As the beta turnover cannot be computed for islands containing a single sampling plot, 1-plot islands were removed from the data (remaining islands = 13). Next, we subsampled the total dataset to obtain a uniform sampling regime consisting of two plots per island. As before, we repeated the sub-sampling process until all possible combinations of 2-plot subsets across the islands in the study system were generated (# subsets = 972). We calculated the beta soundscape turnover by dividing the gamma soundscape richness (pooled richness across 2 plots) by the alpha soundscape richness (mean richness across 2 plots). Finally, we quantified the relationship between the beta soundscape turnover and island area by fitting the power-law model to the total of all two-plots-per-island subsets.

## 3. Results

### 3.1. The effect of patch-scale versus landscape-scape habitat amount on soundscape richness

The partial regression plots show that, when the isolation predictor variable is included in the model, the area predictor is still able to explain a significant proportion of the variation (R^2^ = 0.44; Fig. 2A-left), showing a strong positive relationship between the island size (log_10_) and the unrarefied gamma soundscape richness (log_10_). Conversely, when the island size predictor is included in the model, the isolation variable does not seem to explain any meaningful additional variation (R^2^ = 0.04; Fig. 2A-right). Based on the conditioning scatterplot (Fig. 2B), there appears to be an interaction effect between the island size and isolation on the unrarefied gamma soundscape richness, with isolation acting as a modulator variable. As island isolation increases, the effect of island size on the unrarefied gamma soundscape richness becomes shallower and turns negative at the largest isolation class.

**Figure 2:**
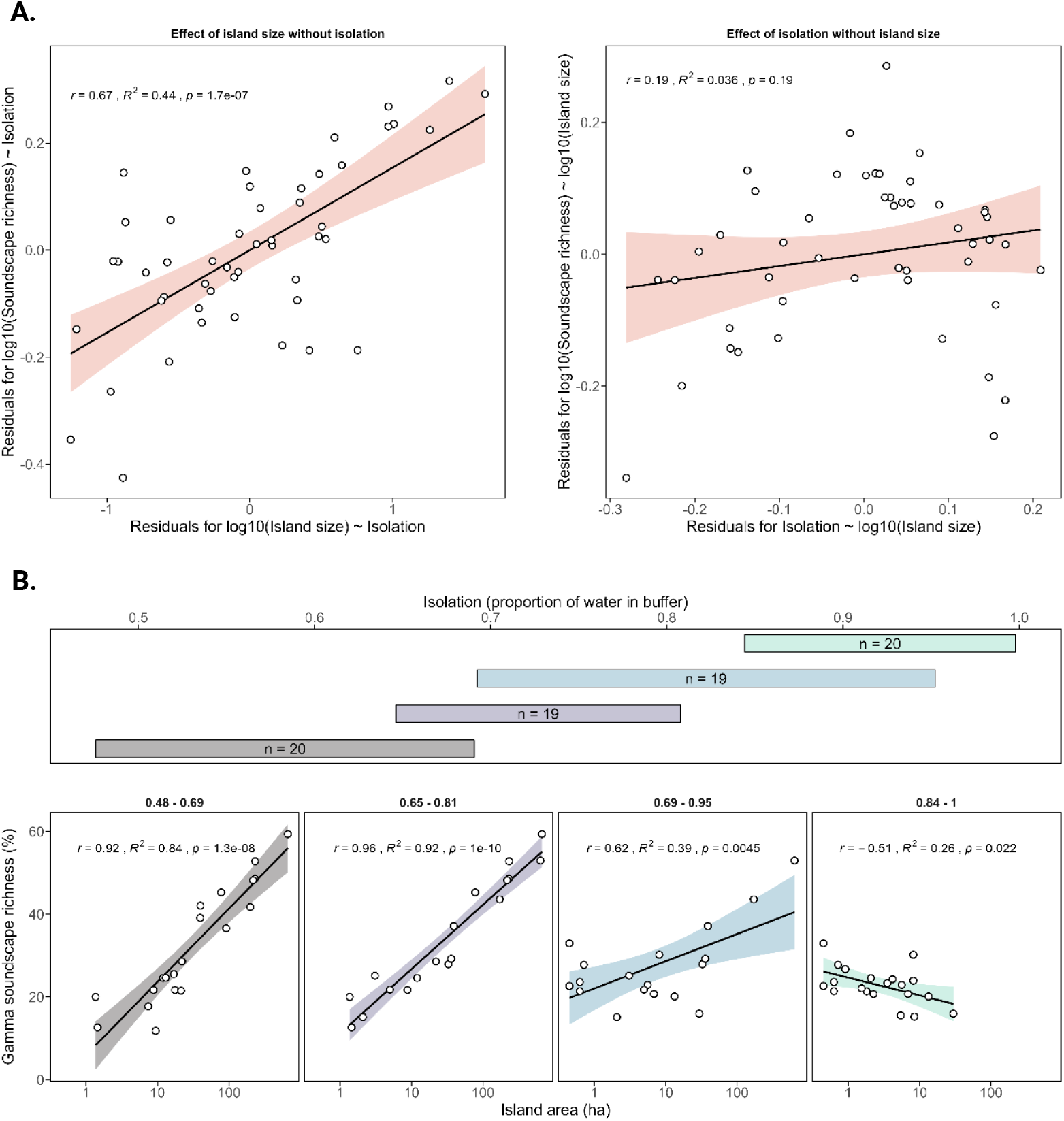
A. Partial regression plots showing the relationship between each predictor variable (left: island size; right: isolation) and the unrarefied soundscape richness while taking into account the variation already accounted for by the other variable; B. A conditioning scatterplot showing the relationship between the unrarefied gamma soundscape richness and island size (log_10_ scale) conditioned on the isolation variable, which is divided into four classes of approximately equal size with 50% overlap between neighbouring classes.

This observation of an interaction effect is confirmed by the model selection. Based on the small sample size corrected Akaike’s Information Criterion and adjusted R^2^-value (AICc = −79.1; R^2^_adj_ = 0.64), the most parsimonious model (model 1) describes the change in log_10_-transformed unrarefied gamma soundscape richness as a function of log_10_-transformed island area, isolation and a log_10_-transformed island size*isolation interaction term (Table 1). The model indicates a highly significant strong negative effect (interaction term: −0.73) of the island isolation on the relationship between the island size and unrarefied gamma soundscape richness. The model abides by most underlying model assumptions, however, the residuals may display a slight deviation from the normal distribution (Table S2; Fig. S7).

**Table 1:**
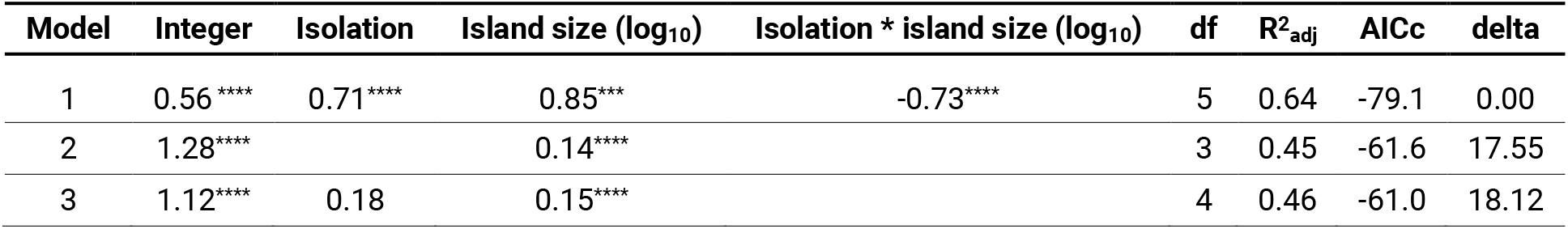
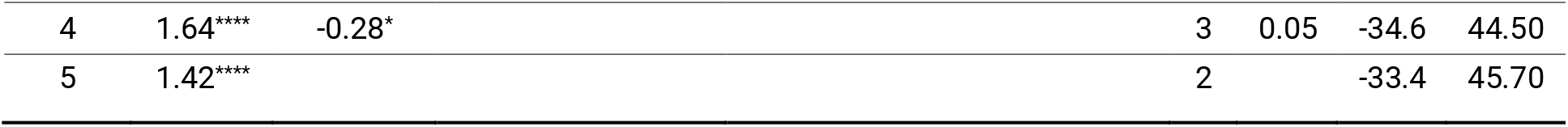
The output of five models compared using an information-theoretic approach. Parameter significance codes: **** = p < 0.001; *** = p < 0.01; ** = p < 0.05; * = p < 0.1;” = p > 0.01.

**Table 2:**
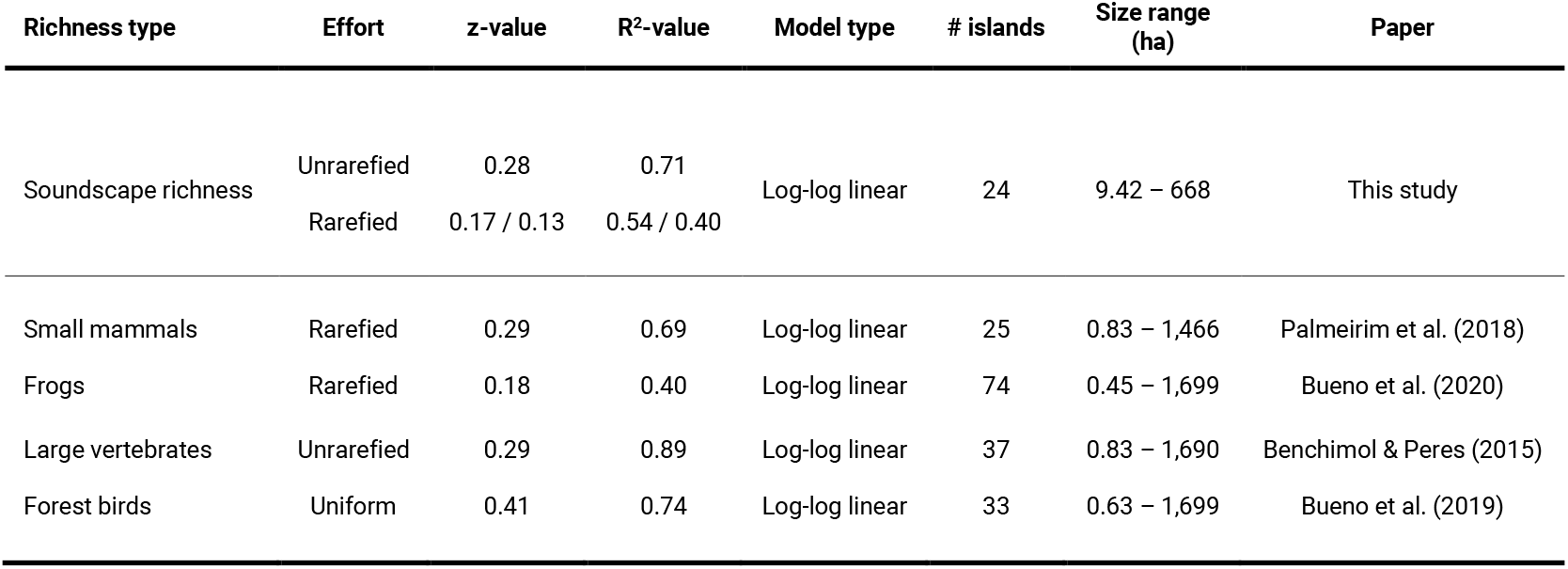
A table displaying the z-(slope) and R2-values for log-log linear power law models fitted to various richness types at the Balbina Hydroelectric Reservoir. Since semi log-linear models were reported in the literature for frogs, for comparability, the raw data presented in the paper’s supplementary material was used to construct log-log linear power models retroactively. For forest birds, no rarefaction was performed since the sampling design had a uniform sampling effort across islands in the study.

| Richness type | Effort | z-value | $R^2$-value | Model type | # islands | Size range (ha) | Paper |
| --- | --- | --- | --- | --- | --- | --- | --- |
| Soundscape richness | Unrarefied | 0.28 | 0.71 | Log-log linear | 24 | 9.42 – 668 | This study |
|  | Rarefied | 0.17 / 0.13 | 0.54 / 0.40 |  |  |  |  |
| Small mammals | Rarefied | 0.29 | 0.69 | Log-log linear | 25 | 0.83 – 1,466 | Palmeirim et al. (2018) |
| Frogs | Rarefied | 0.18 | 0.40 | Log-log linear | 74 | 0.45 – 1,699 | Bueno et al. (2020) |
| Large vertebrates | Unrarefied | 0.29 | 0.89 | Log-log linear | 37 | 0.83 – 1,690 | Benchimol & Peres (2015) |
| Forest birds | Uniform | 0.41 | 0.74 | Log-log linear | 33 | 0.63 – 1,699 | Bueno et al. (2019) |

### 3.2. The ecological mechanisms underlying ISARs

The results from the piecewise regression model selection procedure indicated comparable support for the continuous one-threshold and left-horizontal one-threshold models with threshold values at 9.40 and 12.68 ha respectively, confirming the presence of a small island effect (see supplementary material S5; Fig. S9). As we were interested in the effect of island size on soundscape richness patterns and the mechanisms underlying it, we used the smallest of both threshold values (threshold = 9.40 ha) as the cut-off for small islands in our study. As such, islands smaller than 9.40 ha were excluded from all subsequent analyses (44 plots on 24 islands retained). After small-island removal from the dataset, the power-law model showed an improved positive relationship between the unrarefied gamma soundscape richness and island area in log-log space (Fig. 3A; R^2^_adj_ = 0.71; z-value = 0.28; log_10_c = 1.03) compared to the full dataset (Fig. 3A; R^2^_adj_ = 0.45; z-value = 0.14; log_10_c = 1.28). Though slightly diminished in strength, this relationship persisted when accounting for unequal sampling effort using both temporal-effort-based rarefaction (Fig. 3B-1; R^2^_adj_ = 0.54; z-value = 0.17; log_10_c = 1.15) or plot-based rarefaction (Fig. 3B-2; R^2^_adj_ = 0.40; z-value = 0.13; log_10_c = 1.20).

**Figure 3:**
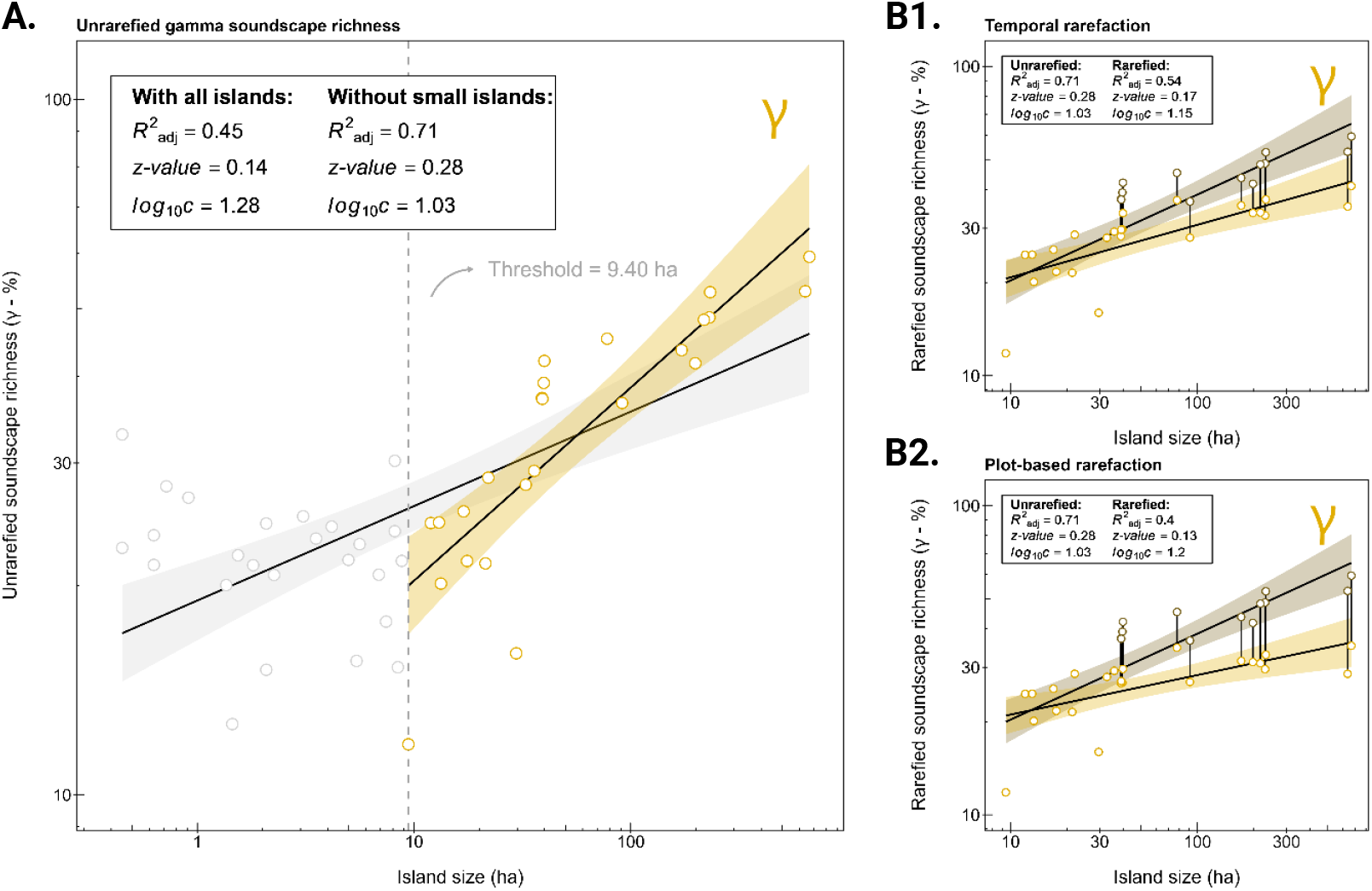
**A**. A scatterplot displaying the relationship between the unrarefied gamma soundscape richness (log_10_) and island size (log_10_) for all islands (n = 49; light grey) and for islands above the 9.40 ha small-island threshold (n = 24; yellow). **B1**. A scatterplot displaying the relationship between the unrarefied (brown) and rarefied (yellow) gamma soundscape richness (log_10_), and island size (log_10_), using temporal effort-based rarefaction (5 sampling days / island) for the islands larger than 9.40 ha. **B2**. A scatterplot displaying the relationship between the unrarefied (brown) and rarefied (yellow) gamma soundscape richness (log_10_), and island size (log_10_), using plot-based rarefaction (1 plot / island) for the islands larger than 9.40 ha. For islands with a single plot, the unrarefied and rarefied values are equal, and thus only the rarefied values (yellow) are displayed.

At the plot-scale, the power-law model applied to all 1-plot subsets showed a positive relationship between the alpha soundscape richness and island area in log-log space (Fig. 4A; adjusted R^2^ = 0.39; z-value = 0.13; log_10_c = 1.20). Conversely, the beta soundscape turnover had no relationship with the island area (Fig. 4B; adjusted R^2^ = 0.00; z-value = 0.00; log_10_c = 0.14).

**Figure 4:**
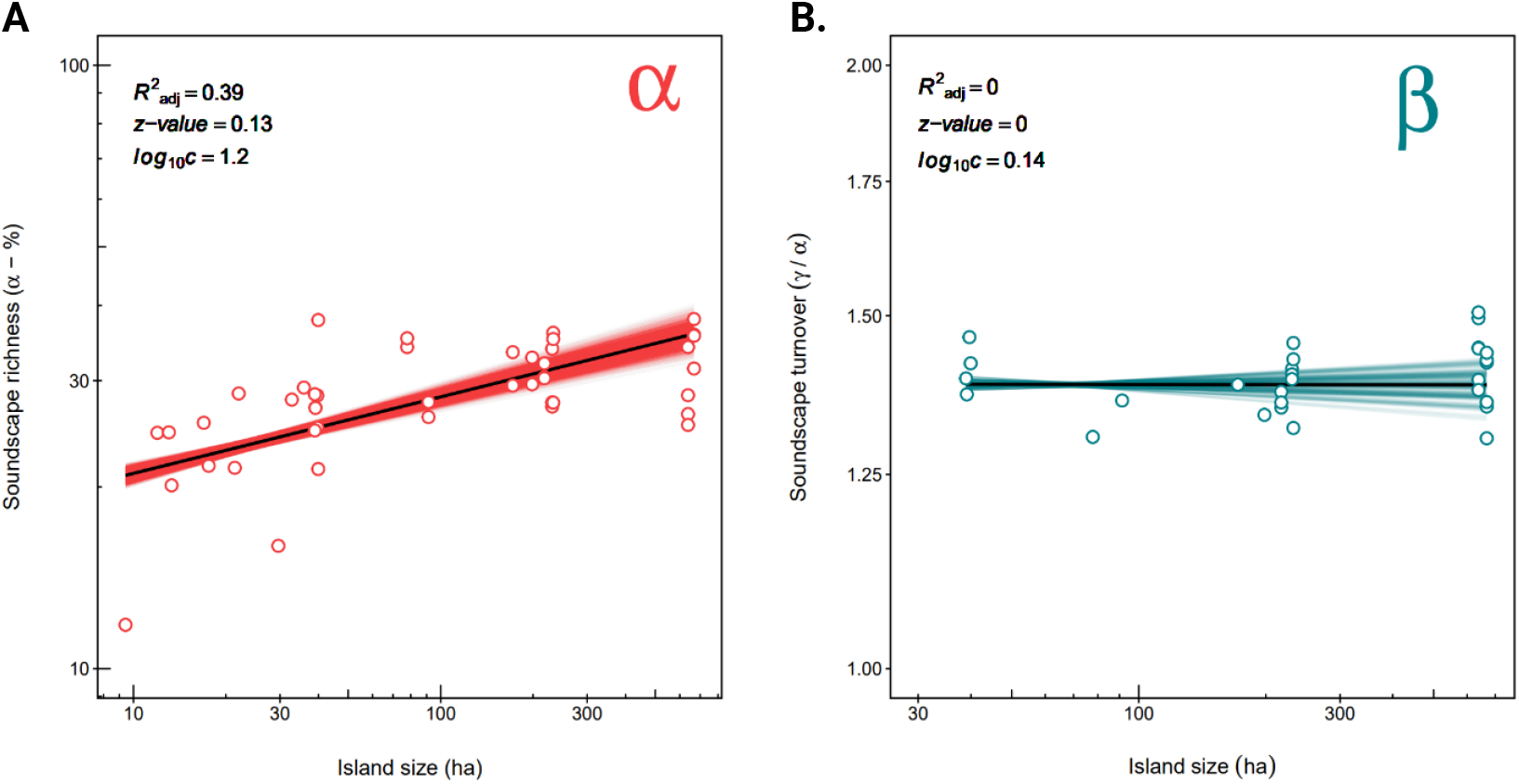
**A**. A scatterplot displaying the relationship between the plot-scale alpha soundscape richness and island area in log-log space (log_10_ scale). **B**. A scatterplot displaying the relationship between the beta soundscape turnover and the island area in log-log space (log_10_ scale). For both plots, the solid black line represents a linear regression fitted to all the data combined (all subset combinations) and the colored lines represent linear regressions fitted to each individual subset (red: 1-plot-per-island subsets; blue: 2-plot-per-island subsets). Due to the large number of subsets for the alpha soundscape richness, approx. 10% of subsets were selected for plotting.

## 1. Discussion

In insular systems, species richness is influenced by island size and isolation, a well-known biogeographic pattern described by the theory of island biogeography. Here, we extended this *ecological law* to the realm of eco-acoustics, testing the relative importance of island size (measured as the patch-scale habitat amount) and island isolation (measured as the inverse of the landscape-scale habitat amount) as descriptors of the spectro-temporal richness of acoustic traits emanating from the landscape, or soundscape richness. Moreover, we decomposed the soundscape richness at sub-island scales and assessed its relationship with island size to gain insight into the ecological mechanisms that may drive the observed patterns in the spatial variation of acoustic trait diversity.

Overall, our best-fit model provides support for a strong positive effect of island size on the unrarefied gamma soundscape richness. Moreover, the presence of a strong negative interaction term in the model suggests the strength of this soundscape richness - island size relationship decreases with an increasing degree of island isolation (Fig. 2B; Table 1). This observed negative interaction effect contrasts with what is expected under the theory of island biogeography (MacArthur and Wilson 1967; Kadmon and Allouche 2007) and what has previously been shown at the BHR (Storck-Tonon and Peres 2017), where the decreasing immigration rates associated with increasing isolation are expected to steepen the ISAR curve. Yet, our island size and isolation predictor variables were negatively correlated, meaning our study system is lacking data on highly connected small islands and highly isolation large islands, and thus, the interaction analysis is comparing different parts of the island size range for different degrees of isolation. For instance, for the lowest levels of isolation, most islands exceeded 10 ha, whereas for the highest levels of isolation, most islands were below 10 ha in size. As we demonstrate a small-island effect later on, with a horizontal or slightly negative ISAR-slope for small islands, it is plausible that the flattening of the ISAR slope for highly isolated (and small) islands, and thus the negative interaction term, is the consequence of stochastic effects obscuring ISAR patterns on small islands, or vice versa. As such, the limitations of the study area and sampling design render it difficult to make inference about the validity of the observed interaction effect.

When we disregard any potential interaction effect between the island size and isolation, we find that a model containing only island size as a predictor best described the variation in the unrarefied gamma soundscape richness (Table 1; model 2; R^2^ _adj_= 0.44; z-value = 0.13; log_10_c = 1.28). Contrary to what is expected under the IBT, we found no effect of island isolation on the unrarefied gamma soundscape richness. Moreover, as we measured the island isolation as the inverse of the landscape-scale habitat amount, the absence of an isolation effect suggests that the sampling effect underlying the habitat amount hypothesis (Fahrig 2013) is not the driving mechanism behind the soundscape richness gradient. The importance of island-versus landscape-scale factors in governing species richness regulation in insular systems is dependent on the degree of ‘*islandness’* of the habitat patches, which in turn depends on the matrix permeability and species dispersal abilities (Bueno and Peres 2019). The lack of a landscape-scale habitat amount effect (HAH) is expected at the BHR, as the habitat patches are separated by an inhospitable open-water matrix that is largely prohibitive to between-patch dispersal for many taxonomic groups (Bueno et al. 2020). Other than euglosine bees and ant-myrmecophyte networks (Storck-Tonon and Peres 2017; Emer et al. 2013), the absence of isolation effects has previously been described for birds (Aurélio-Silva et al. 2016), large vertebrates (Benchimol and Peres 2015b), lizards (Palmeirim et al. 2017) and harvestmen (Tourinho et al. 2020) at the Balbina Hydroelectric Reservoir.

Although ISAR patterns have been demonstrated numerously for taxonomic-richness-based studies and are expected from several theoretical predictions (MacArthur and Wilson 1967; Connor and McCoy 1979), to our knowledge, this represents the first empirical evidence for a positive relationship between the island-scale (gamma) soundscape richness and island size, which we term a *SoundScape-Area Relationship* (SSAR). This term was previously coined in de Camargo et al. (2019), however, these authors derived traditional taxonomic richness data from sound files and thus captured a conventional species-area relationship using acoustic methods, which does not fully justify the use of a novel term. Yet, positive ISAR patterns can arise through a range of mechanisms, including sampling effects (sampling artefacts and passive sampling), disproportionate effects and heterogeneity effects. We disentangled which of these mechanisms generated the SSAR by dissecting the soundscape richness at various spatial scales.

Prior to assessing the mechanisms underlying the observed SSAR, we tested for the presence of a small-island threshold, below which island size has no effect on the soundscape richness. The piecewise regression analysis revealed a comparable fit for the continuous one-threshold and left-horizontal one-threshold models, with respective thresholds at 9.40 and 12.68 ha, confirming the presence of a small-island effect. Small islands at the BHR are affected by severe edge effects, including windfalls and ephemeral wildfires, resulting in a strong reduction in structural habitat complexity and resource availability (Benchimol and Peres 2015a). The stochasticity associated with these edge effects can obscure the effect of island area on the soundscape richness, leading to the breakdown of the ISARs at small spatial scales. Indeed, the existence of SIEs has previously been suggested at the Balbina Hydroelectric Reservoir for several sound-producing taxonomic groups. For instance, Bueno et al. (2020) found that, for anurans, ISARs had weak inferential power below 100 ha. Similarly, both large vertebrates and understory birds had much shallower or non-existent ISARs below 10 ha (Benchimol and Peres 2015b; Bueno and Peres 2019). When disregarding islands smaller than 9.40 ha, the relationship between the unrarefied gamma soundscape richness and island area displayed an improved fit (adjusted R^2^ = 0.71; z-value = 0.28; log_10_c = 1.03).

Next, we investigated the null hypothesis that ISARs are simply the result of sampling effects by equalizing the sampling effort using two rarefaction approaches. Though slightly weakened, we found a positive relationship between the soundscape richness and island area for both the temporal (Fig. 3B-1; R^2^_adj_ = 0.54; z-value = 0.17; log_10_c = 1.15) and plot-based (Fig. 3B-2; R^2^_adj_ = 0.40; z-value = 0.13; log_10_c = 1.20) rarefaction, suggesting the relationship is driven by underlying mechanisms beyond sampling artefacts. As expected, the slope and R^2^-values of the soundscape-area relationship are slightly weaker than for most taxonomic-richness-based studies reported in literature for the BHR (Table 1). This is expected, as acoustic indices can be sensitive to non-target sounds, including abiotic sounds such as rain, wind or vegetation (Gasc et al. 2015), thus introducing variability into the model. Moreover, as demonstrated in Bueno et al. (2020), the strength of the effect and explanatory power of SAR-models are sensitive to the range of islands sizes under consideration, with both z- and R^2^-values increasing as larger islands are included. The upper range of island sizes included in this study (668 ha) was smaller than other SAR-studies in the area (approx. 1,500 ha), thus potentially depressing the observed slope and R^2^-values of the log-log linear model.

As passive sampling effects are likely always operating in the background (Chase et al. 2019), we tested whether other SAR-generating mechanisms were active by assessing the relationship between island size, and the alpha soundscape richness and beta soundscape turnover (Schoereder et al. 2004). The local alpha soundscape richness displayed a positive relationship with island area (Fig. 4A; adjusted R^2^ = 0.39; z-value = 0.13; log_10_c = 1.20), indicating that SSARs were at least partly generated by disproportionate (Schoereder et al. 2004) or heterogeneity effects (through spillover effects - see Giladi et al. 2014). The absence of a relationship between the beta soundscape turnover and island size (Fig. 4B; adjusted R^2^ = 0.00; z-value = 0.00; log_10_c = 0.14) rules out heterogeneity effects, validating disproportionate effects as the main mechanism underlying the SSAR. This suggests that remnant size affects the processes that regulate soundscape richness at a local scale. A meta-analysis of plant ISARs found that approximately 40% of studies provided evidence for a positive relationship between the alpha richness and island size (Giladi et al. 2014). Moreover, Chase et al. (2019) found that disproportionate effects were an important SAR-generating mechanism for grasshoppers and lizards, but not for shrubs. Here, the authors posited that the importance of disproportionate effects versus sampling effects is dictated by the type of matrix and group under investigation. For communities isolated by more hostile matrices, or groups with a lower dispersal ability, local processes likely outweigh regional sampling effects, leading to disproportionate effects. Indeed, the hostile water matrix at the BHR could help explain the importance of disproportionate effects in generating the observed SSARs.

In addition to shedding light on the ecological mechanisms that modulate soundscape richness along an area gradient, the study design also allows us to posit which mechanisms generate acoustic trait diversity in the first place. The positive relationship between the island size and plot-scale alpha soundscape richness, and the absence of a relationship with the beta soundscape turnover, suggests that the landscape-scale richness of acoustic traits is unlikely to be governed by environmental filtering for optimal sound propagation (acoustic adaptation hypothesis), as in this case we would expect the composition of acoustic traits to be uniquely adapted to each habitat and showcase minimal overlap (Mullet et al. 2017). Instead, we deem it more likely that the soundscape richness is primarily influenced by the diversity of sound-producing organisms, as posited by the acoustic niche hypothesis (Krause 1993). Indeed, a strong positive relationship between the soundscape richness and taxonomic richness of soniferous organisms (frogs, birds and primates) has previously been show at the Balbina Hydroelectric Reservoir (Luypaert et al. 2022).

The ANH states that, over evolutionary timescales, undisturbed ecosystems acquire an equilibrium between sounds in the landscape, resulting in soundscapes with high spectro-temporal complexity and signal diversity, and minimal overlap (Eldridge et al. 2016; Krause 1993; Pijanowski et al. 2011a; Pijanowski et al. 2011b). In fragmented landscapes such as the one in this study, this equilibrium is disturbed and locally-adapted species are lost from the ecosystem as island size decreases. Previous work at the Balbina Hydroelectric Reservoir has shown that the loss of species along the island area gradient occurs non-randomly, as forest-bound specialist species tend to be lost or replaced with generalist species, leading to functional impoverishment (Palmeirim et al. 2017). In the context of the ANH, it is plausible that acoustically optimized species are lost from their ecosystem and replaced with generalists that are not adapted to the specific acoustic environment they inhabit, leading to overlap in acoustic signals, readily detectable gaps in the soundscape, and ultimately, a lower soundscape richness for more defaunated islands.

Although rarely assessed, previous work in fragmented Amazonian landscapes has demonstrated that also the abundance of species decreases with island size. For instance, for large rainforest vertebrates, the abundance of nearly all species decreased with habitat patch or island size (Michalski and Peres 2007; Benchimol and Peres 2021). Moreover, island size also affected the maximum operational group size of several vocal social species such as primate species and trumpeters, with suppressed group sizes on small islands (Benchimol and Peres 2021). It is likely that these abundance-area relationships contribute to the observed soundscape-area relationship to some extent. For social species, both overall calling rates (Payne et al. 2003) and individual calling rates (Radford and Ridley 2008; Fernandez et al. 2017) have been shown to correlate positively with group size. Moreover, it is thought that animals that live in more complex social communities exhibit more complex vocal repertoires, as individuals have to navigate more vocal interactions within and between social groups (Teixeira et al. 2019). The simplified acoustic environment resulting from defaunation on smaller islands may lead species to exhibit less elaborate vocal repertoires with lower spectro-temporal complexity and signal diversity. Moreover, for vocalizations associated with competition for resources, lower abundance of conspecifics on smaller islands may lead to a reduction in calling rates due to competitive release (Radford and Ridley 2008). As acoustic indices have previously been shown to correlate positively with species abundance (Boelman et al. 2007; Fuller et al. 2015; Bradfer-Lawrence et al. 2020), it is plausible that these factors also influence the soundscape richness metric used in this study.

## Supplementary material 1: Data collection

### 1.1. Collection of acoustic data

#### The study system

Hydroelectric reservoirs represent an excellent study system to assess the spatial scaling of biodiversity. On the one hand, they constitute a rapidly emerging threat to Neotropical rainforest ecosystems (Finer and Jenkins 2012; Emer et al. 2013; Fearnside 2006). On the other hand, hydroelectric reservoirs are considered a perfect experimental system to study the effects of island area while controlling for confounding effects. For instance, they allow us to capitalise on the fact that all patches were formed simultaneously due to a single disturbance event. Moreover, they have a uniform and largely untraversable matrix, a spatial scale comparable to terrestrial patches and were formed recent enough so that evolution and species adaptation have yet to take effect.

This study was conducted at the Balbina Hydroelectric Reservoir (BHR) in Brazilian Amazonia (1°40′S, 59°40′W; Fig. 1), one of the largest hydroelectric reservoirs on Earth (Bueno et al. 2020). The reservoir was formed when a tributary of the Amazon, the Uatumã River, was dammed in 1987, turning the former hilltops of primary continuous forest into > 3,500 islands spanning an area of approximately 300,000 ha (Bueno and Peres 2019). The artificial tropical rainforest archipelago now contains islands spanning a wide range of sizes, ranging from 0.2 to 4,878 ha (Benchimol and Peres 2015a). The area’s vegetation is characterised by submontane dense ombrophilous (*terra firme*) forest. Moreover, the forest structure of larger islands resembles the continuous forest, with large-seeded and canopy tree species dominating assemblies. Conversely, smaller islands are dominated by pioneer species due to edge effects (Benchimol and Peres 2015a). Finally, virtually all islands lack perennial streams due to the submergence of lowland areas.

##### Data collection

*see Bueno and Peres (2019) for detailed overview of data collection*

Acoustic surveys were conducted between July and December 2015, recording long-duration acoustic data at 151 plots on 74 islands and 4 continuous forest sites. As previously mentioned, the number of plots per area was proportional to the habitat area associated with each plot, and varied between 4-10 for continuous forest sites and 1-7 for islands. At each plot, a passive acoustic sensor (an LG smartphone enclosed in a waterproof case linked to an external omnidirectional microphone) was attached to a tree trunk at 1.5m height and set to record the soundscape for 1 minute every 5 minutes for 4-10 days at a sampling rate of 44,100 Hz using the ARBIMON Touch application (ARBIMON, https://arbimon.rfcx.org/).

#### Site selection

Several islands and sampling plots were removed from this study: (i) mainland sites; (ii) riparian habitats; (iii) plots with microphone failure; and (iv) plots with overly noisy recordings. Due to the removal of these plots from the study data, some islands deviated from the proportional sampling regime. We rectified this by retroactively removing islands that deviated from the proportional sampling regime (r = 0.87; R^2^ = 0.76; p < 0.001; Fig. S1). Ultimately, we retained 69 plots on 49 islands for further analysis (Table S1).

**Figure S1:**
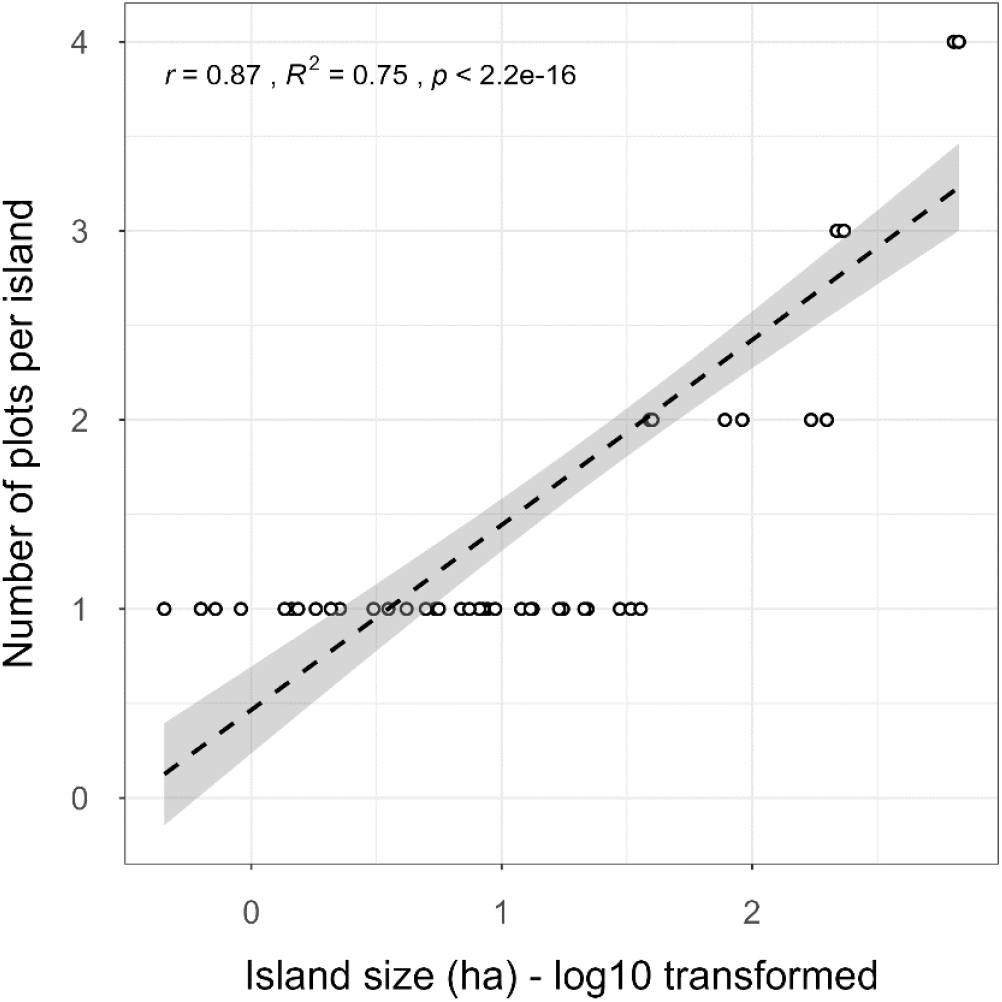
A scatterplot with regression line showing the proportional relationship between the island size (log_10_ transformed) and the number of acoustic sampling plots per island (r = 0.87; R^2^ = 0.75; p < 0.005) for the final set of islands used in the study.

**Table S1:**
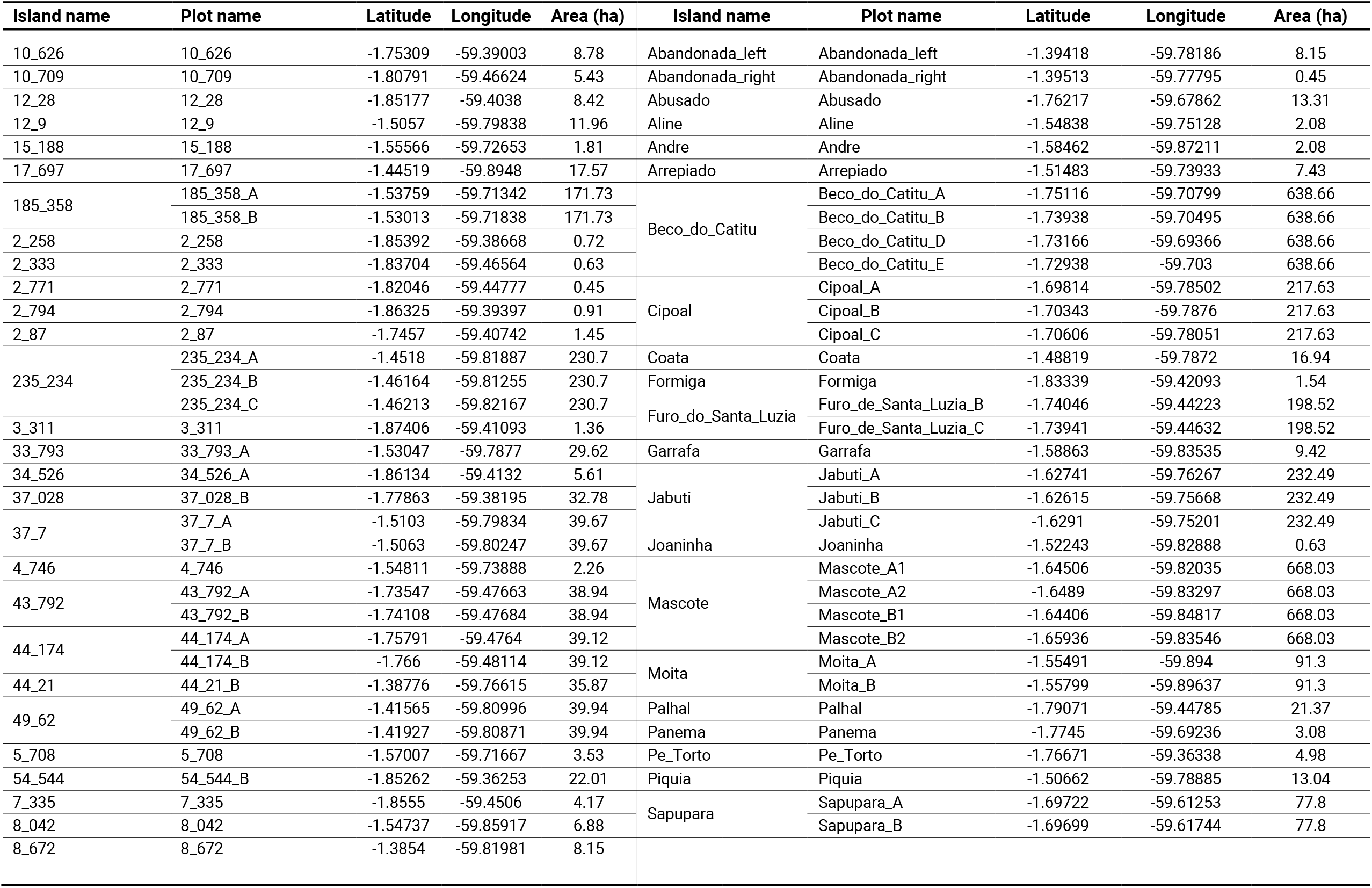
An overview of the sites included in the study

## Supplementary material 2: Soundscape richness

To quantify the richness of acoustic traits emanating from the landscape for each island in the study, we employed the analytical pipeline outlined in Luypaert et al. (2022) to calculate the *soundscape richness* acoustic index. As soundscape diversity metrics aim to capture ecological patterns without the need to isolate and identify species’ vocalizations from sound files, they generally lack a unified unit of diversity measurement (e.g. species). To overcome this, the pipeline groups sounds by their shared spectro-temporal properties in the 24h trait space in which species produce sound, better known as ‘Operational Sounds Units’ or OSUs. Using an incidence-approach, the relative abundance of these OSUs is quantified throughout the recording period and soundscape diversity metrics are calculated using the analytical framework of Hill numbers.

To assess potential patterns of soundscape richness in our insular system and elucidate which underlying mechanisms might drive these, we quantified the soundscape richness using an adapted version of the multi-scale and multi-metric framework outlined in Chase et al. (2019). We analysed the spatial biodiversity scaling using four soundscape metrics at various spatial scales: (i) the unrarefied island-wide gamma soundscape richness; (ii) the rarefied island-wide gamma soundscape richness; (iii) the local plot-scale alpha soundscape richness; and (iv) a beta soundscape turnover metric between plots per island.

### Gamma soundscape richness

#### Unrarefied gamma soundscape richness

To assess the relative importance of island size (patch-scale habitat amount) versus island isolation (inverse of landscape-scale habitat amount), and uncover the shape of the island soundscape-area relationship (SSAR), firstly, we quantified the unrarefied island-wide gamma soundscape richness. To do so, we pooled the OSU-by-sample incidence matrices across all plots per island. Per example, for an island consisting of 4 sampling plots with five sampling days each, pooling the OSU-by-sample incidence matrices across all plots (each matrix consisting of 5 columns of OSU detection (1) / non-detection (0) data for that sample) would result in a pooled matrix consisting of 20 columns (soundscape samples) containing OSU incidence data. Using the pooled OSU-by-sample incidence matrix per island, we quantified the unrarefied gamma soundscape richness by counting how many unique OSUs were detected across all soundscape samples.

#### Rarefied gamma soundscape richness

As we employed a proportional sampling scheme, for which larger islands were sampled more intensely, we might expect a positive relationship between the unrarefied gamma soundscape richness and our predictor variables because of sampling effects (*sampling artefacts*). To account for this unequal sampling effort between islands, we adopted a rarefaction procedure to calculate the rarefied gamma soundscape richness per island. The framework outlined in Chase et al. (2019) employs an individual-based rarefaction framework to equalize the sampling effort between islands. However, due to the nature of the soundscape richness metric used here, for which we renounced the isolation and identification of individual vocalizations from sound files, this type of individual-based abundance data is not available. Instead, the soundscape richness metric was calculated using sampling-unit-based incidence data. As such, to equalize the sampling effort among island, we employed a *sample-based* rarefaction procedure.

For our workflow, what exactly constitutes a sample of the soundscape in the OSU-by-sample incidence matrix is dependent on the scale at which we regard the soundscape. At a plot scale, each 24h period in the acoustic survey represents a sample of the soundscape for which we determine the detection (1) / non-detection (0) of OSUs. As such, when performing sample-based rarefaction, the temporal sampling effort (number of 24h sampling periods in the acoustic survey) is equalized.

However, when regarding the soundscape richness on an island-scale, we have multiple sampling plots per island, each with its own OSU-by-sample incidence matrix. When rarefying the sampling effort at this scale, what constitutes a sample of the soundscape can be viewed in one of two ways. If we don’t take the spatial heterogeneity between plots into account and assume that OSUs are distributed randomly across the island, we can pool the OSU-by-sample incidence matrices across all plots on the island into one island-scale OSU-by-sample incidence matrix and equalize the sampling effort between islands using the number of 24h soundscape samples. However, if habitat heterogeneity influences the presence and distribution of OSUs across the island, we expect that an increase in the number of plots per island will inflate the number of detected OSUs. In this case, rather than treating each 24h sampling period in the acoustic survey as a soundscape sample, we should treat each plot as a sample of the soundscape and rarefy the sampling effort using the number of plots per island. In this study, to account for the potential confounding influence of sampling artefacts on the soundscape-area relationship, we rarefied the island-wide gamma soundscape richness using both the temporal- and plot-based rarefaction procedures.

For the temporal rarefaction, we pooled the OSU-by-sample incidence matrices across all plots per island and rarefied the sampling effort between islands to 5 sampling days. For the plot-based rarefaction, we converted each plot’s OSU-by-sample incidence matrix to a vector containing the detection (1) / non-detection (0) of OSUs at that plot across the whole acoustic survey. Then, we constructed a novel island-scale OSU-by-sample incidence matrix with plots as samples. We performed plot-based rarefaction to one plots per island. The results of relationship between island size and the rarefied soundscape richness are presented in the main text (section 3.2).

## Supplementary Material 3: Calculating the isolation variable

To examine the degree to which landscape-scale habitat amount (or island isolation) was informative, we follow the approach outlined in MacDonald et al. (2018). The amount of isolation was quantified for all islands in the study by calculating the proportion of water (1 – proportion of land area) within a range of buffer sizes calculated from the island edge. To determine the optimal scale-of-effect for our isolation variable (Jackson and Fahrig 2015), we considered 40 different buffer sizes, ranging from 50 to 2000 m at 50-m intervals.

The computation of the isolation metric was performed using a combination of the open-source QGIS software (QGIS Association 2022 - version 3.2.2) and the R environment (R Core Team 2022 - version 4.1.2).

### 1.1. Deriving the total land area shapefiles for islands in the study

First, we downloaded a land cover map for Brazil in 2015 (the time at which the study was conducted) using the ‘MapBiomas’ online download tool (collection 6: https://storage.googleapis.com/mapbiomas-public/brasil/collection-6/lclu/coverage/brasil_coverage_2015.tif). Next, the MapBiomas GeoTIFF raster file was uploaded into QGIS and clipped to contain the study landscape (the Balbina Hydroelectric Reservoir). Next, the ‘identify features’ tool was used to sample the raster values corresponding to the water matrix within the Hydroelectric Reservoir (value = 33). We then used the ‘raster calculator’ to binarize the raster values, setting the water matrix values to 0, and all surrounding land area to 1. The binary matrix was then converted to a set of polygon shapefiles using the ‘polygonize’ function. To retain only the land area as polygons and remove the water matrix from the shapefile, the attribute table of the polygonized binary raster data was edited, removing all shapefiles containing zero values. This shapefile was merged with the previously generated ‘island forest area’ shapefile to get the final islands shapefile for the study area. To do so, we used the ‘merge’ and ‘dissolve’ functions, and patched any holes within polygons using the ‘delete holes’ function. A visual inspection of the island shapefiles with an ESRI satellite base map revealed an excellent approximation of island shapes at the Balbina Hydroelectric Reservoir.

### 1.2. Generating the buffer rings around the islands in the study

We further isolated individual islands from the final shapefiles of all island using the ‘*multiparts to singleparts’* function and selected the islands in the study area based on the coordinates of the sampling plots using the ‘select by location’ function. For all islands included in the study area, we generated 40 buffers around each focal island using a range of buffer sizes calculated from the island edge (from 50m to 2000m at 50m intervals). Since we want the buffers to exclude the original island from the buffer area, we used the ‘*multi-ring buffer*’ QGIS plugin function. For each buffer size, we calculated the total area covered by the buffer using the attribute table field calculator. Next, to calculate the area covered by land (island or mainland) within each buffer area, we used the ‘*clip’* function, clipping the buffer shapefiles with the shapefile of all islands. Then, we calculated the total land area within each buffer using the attribute table field calculator. The attribute tables were exported from QGIS and imported into R for further analysis. For each island and buffer size, the proportion of land within the buffer area was calculated by dividing the land area by the total area of the buffer. Finally, the proportion of open-water within the buffer area was calculated (proportion of water = 1 – the proportion of land). In doing so, we obtained a metric of isolation (proportion of water within the surrounding buffer) which ranged from 0 to 1 for 40 different buffer size around each of the islands in this study.

### 1.3. Determining the scale-of-effect

To determine the spatial scale at which the isolation metric attains the strongest relationship with soundscape richness, we calculated the correlation between the unrarefied island-wide gamma soundscape richness and the proportion of water for each of the 40 buffer sizes under investigation.

**Figure S2:**
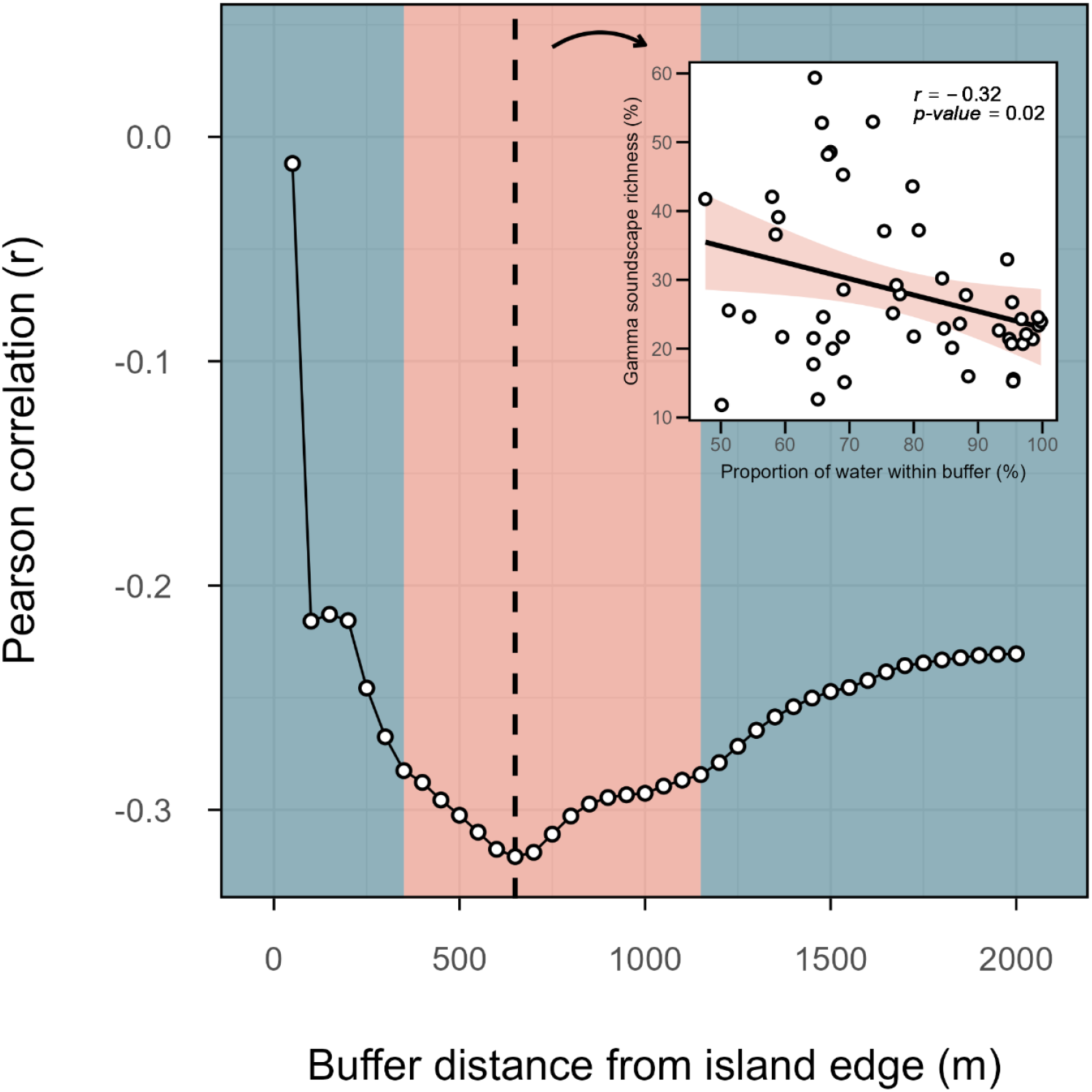
A visual representation of the spatial scale at which the landscape-scale isolation metric derived in this study (proportion of water within the buffer area of each island) attains the highest correlation value with the soundscape richness metric (unrarefied gamma soundscape richness); this is better known as the ‘scale-of-effect’ (Jackson and Fahrig 2015). The blue shading represents buffer distances for which the correlation between soundscape richness and the isolation metric were not significant (p > 0.05), whereas orange shading represents significant correlations. The highest correlation value was reached at a buffer distance of 650 m. The inset plot displays the correlation between the soundscape richness and our isolation metric at the scale-of-effect (650 m).

The highest correlation between soundscape richness and our isolation metric was attained at a scale-of-effect of 650 m (Fig. S2). At this spatial scale, we observed a significant negative correlation between the degree of island isolation and soundscape richness (r = −0.32; p < 0.05).

## Supplementary Material 4: Modelling island size versus isolation

To examine the effect of island size versus island isolation on the unrarefied island-wide gamma soundscape richness, we fitted a series of linear models and used an information theoretic approach for model selection (Burnham and Anderson 2004).

### 4.1. Checking variable distributions

Before modelling the data, we assessed the distribution of the three variables under investigation (unrarefied gamma soundscape richness, island size, and island isolation) using raincloud plots and Quantile-Quantile plots (Fig. S3, S4 and S5). We observed a right-skewed distribution for the gamma soundscape richness and island area (Fig. S3 and S4). To account for this, these variables were log-transformed (log_10_ x).

**Figure S3:**
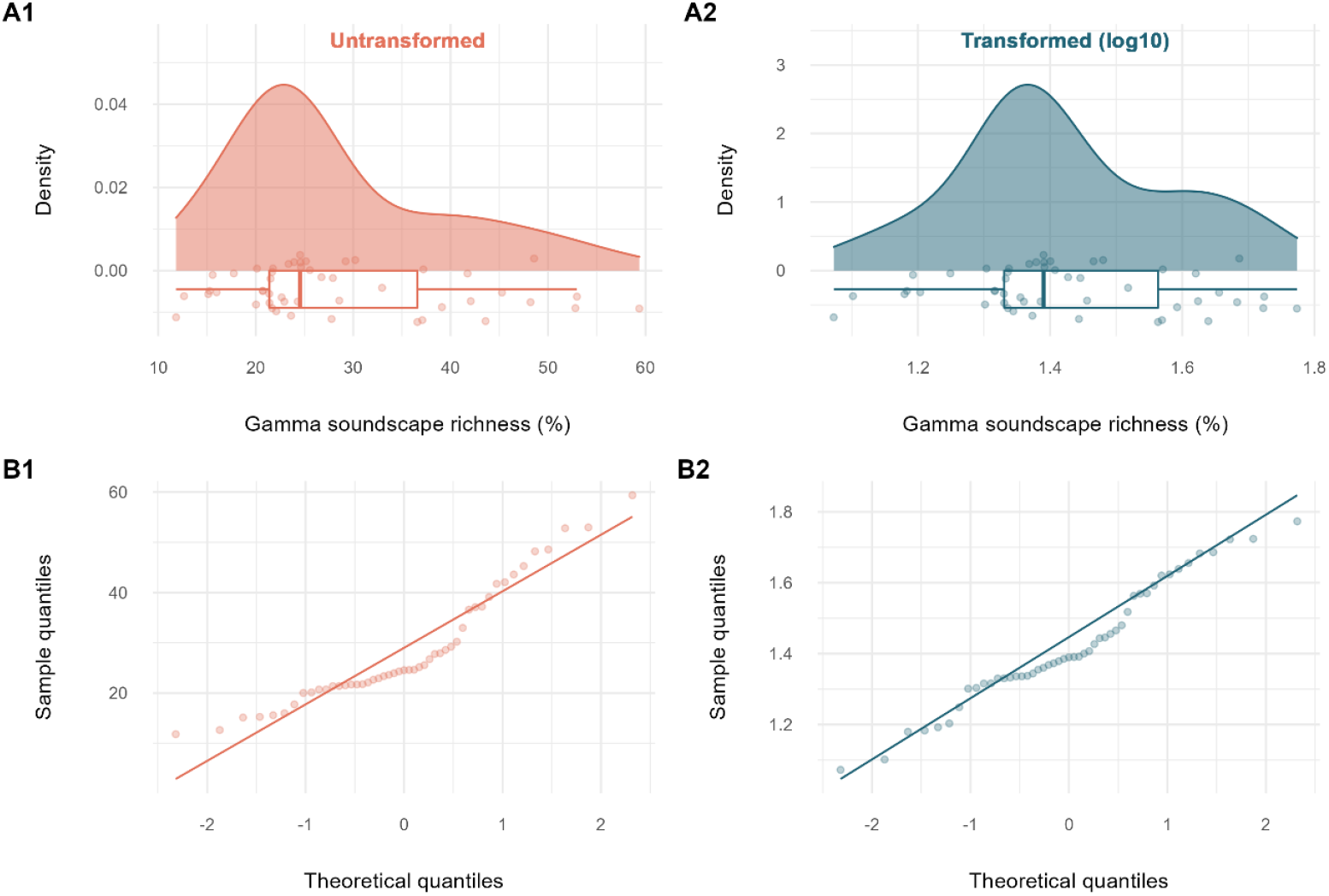
A visual representation of the distribution of unrarefied gamma soundscape richness using (A) density plots and (B) quantile-quantile plots for both untransformed (1; orange) and log_10_-transformed (2; blue) data.

**Figure S4:**
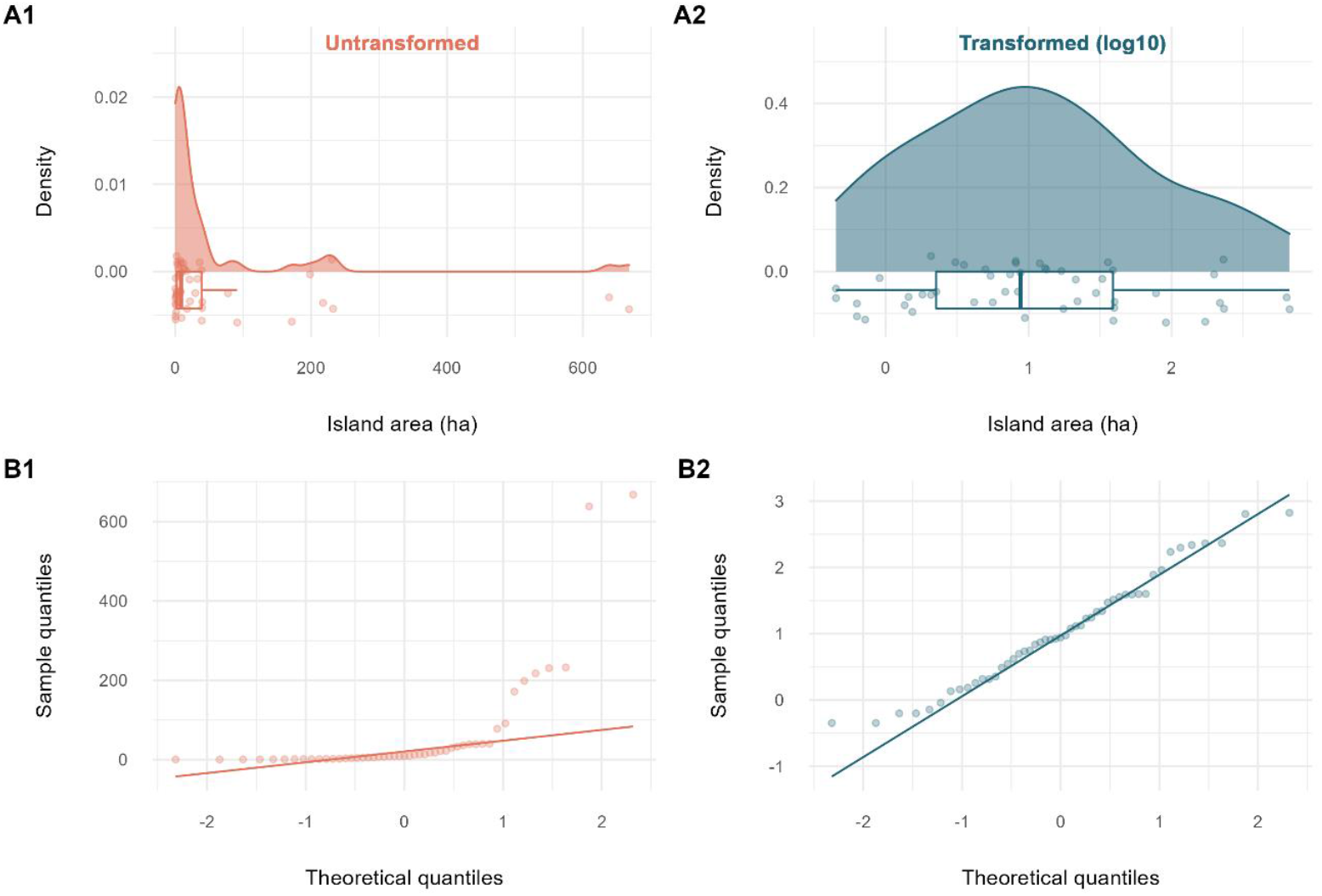
A visual representation of the distribution of the island size variable using (A) density plots and (B) quantile-quantile plots for both untransformed (1; orange) and log_10_-transformed (2; blue) data.

**Figure S5:**
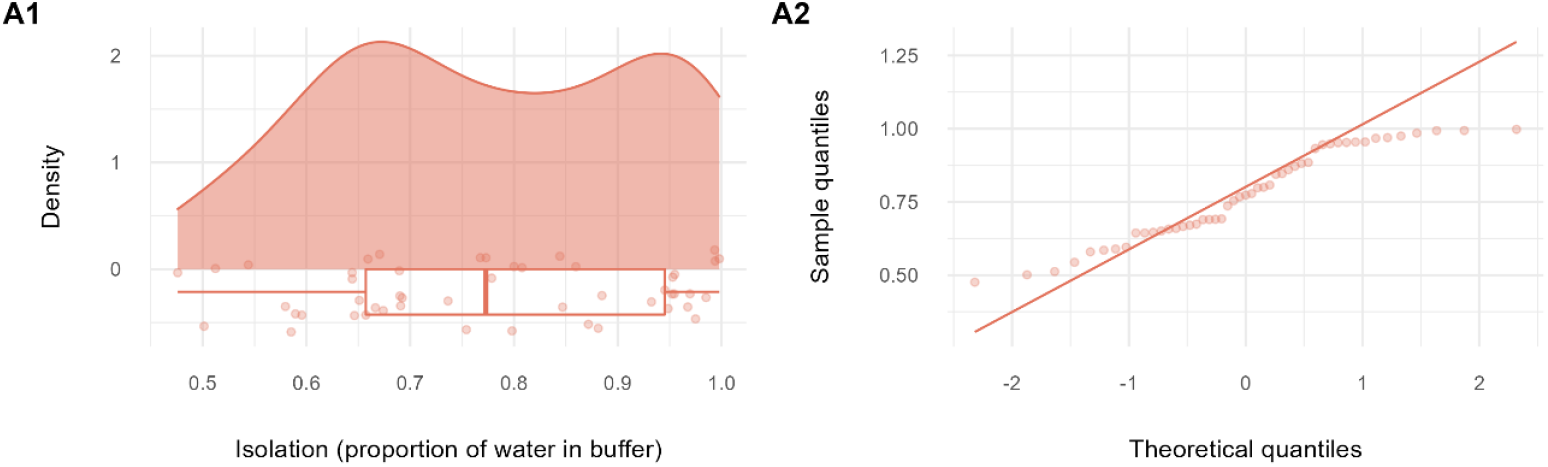
A visual representation of the distribution of the isolation variable (proportion of water within buffer) using (A1) density plots and (A2) quantile-quantile plots.

### 4.2. Exploring the relationship between predictor variables

In fragmented landscapes, there is often a correlation between the island size and isolation, with smaller islands also being more isolated, or larger islands more connected. As this correlation could have consequences for subsequent modelling procedures, first, we explored the relationship between the size and isolation for the islands contained in the study (Fig. S6).

**Figure S6:**
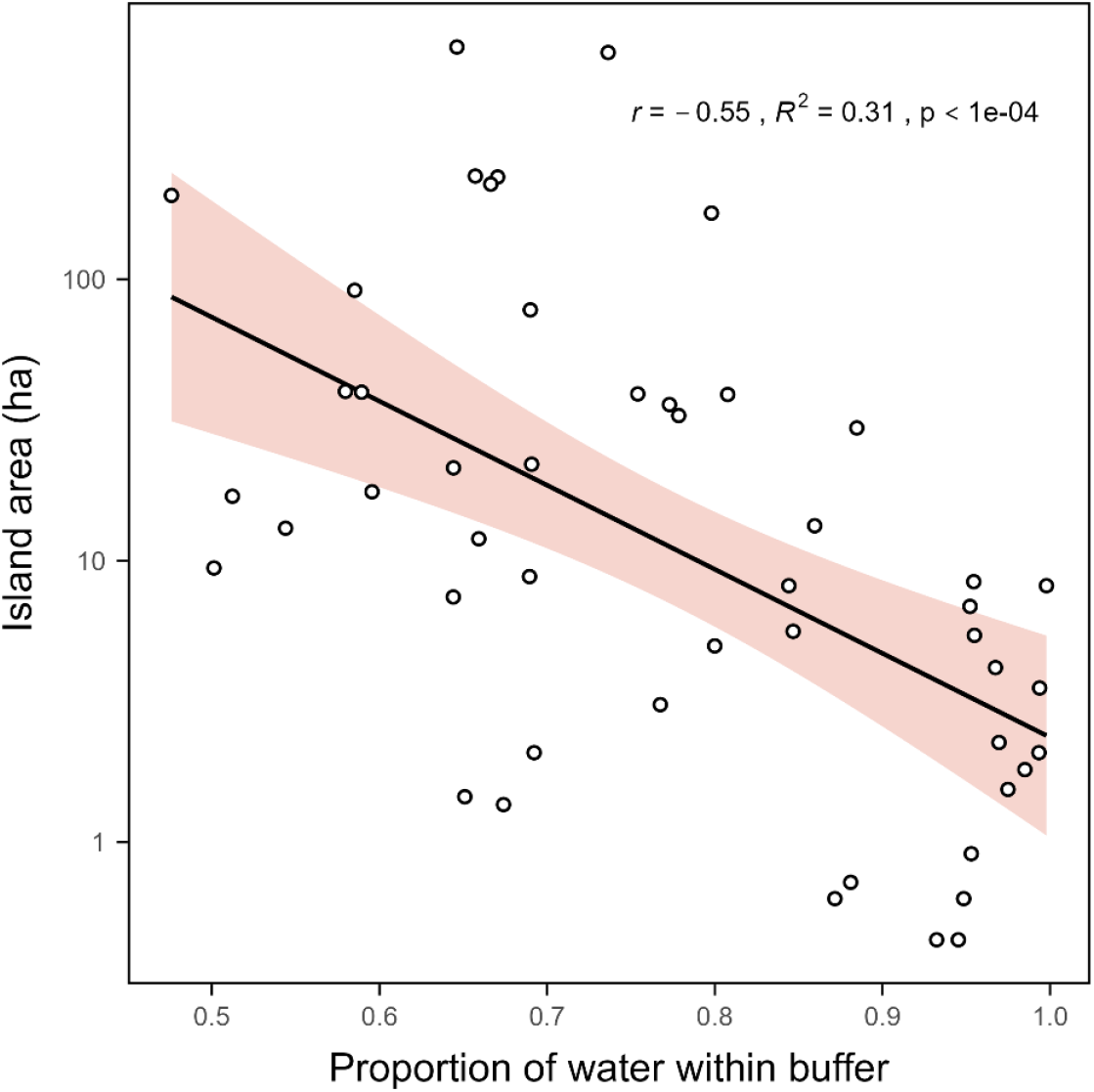
A scatterplot displaying the relationship between the island area (ha – log_10_ scale) and island isolation (proportion of water within 650 m buffer around the island edge).

Indeed, as anticipated, our two predictor variables have a strong and negative correlation, a typical observation in fragmented landscapes. Our study system is missing highly connected small islands and highly isolated large islands. We will keep this in the back of our mind as we continue our analysis.

### 4.3. Exploring the relationship between predictor and response variables using partial regression plots

Before fitting our linear regression models, we visually explored the relationship between the unrarefied gamma soundscape richness, and the island size and isolation. However, when using regressions that involve multiple potential predictor variables, the use of bivariate plots to show the relationship between X~Y may be misleading, as the regression coefficients may change in magnitude and sign when more than one predictor has an influence on the response variable (Moya-Laraño and Corcobado 2008). Instead, we made use of *partial regression plots* (also known as *added variable plots* or *adjusted variable plots*), which allowed us to plot the effects of area and isolation separately while accounting for the variation taken up by the other variable (Fig. 3A).

Say we are interested in the relationship between log_10_(soundscape richness) ~ log_10_(island size), the plot displays the relationship between the residuals of a model between log_10_(soundscape richness) ~ isolation and the residuals of a model between log_10_(island size) ~ isolation. In doing so, the plot shows the relationship of log_10_(island size) on the soundscape richness while eliminating the effect of isolation. Similarly, if we are interested in the relationship between log_10_(soundscape richness) ~ isolation, the plot displays the relationship between the residuals of a model between log_10_(soundscape richness) ~ log_10_(island size) and the residuals of a model between isolation ~ log_10_(island size).

### 4.4. Assessing the presence of an interaction effect between island size and isolation

In addition to visually exploring the relationship between the predictors and the response variable, we were also interested in assessing whether an interaction effect between our continuous predictor variables existed. To test this, we made use of conditioning plot (also known as a co-plot), a type of scatterplot that shows the relationship between two variables when ‘*conditioned*’ on a third variable. In our case, we visualized the potential interaction between the island size and isolation by plotting the relationship between the log_10_(soundscape richness) ~ log_10_(island size) for four classes along the isolation range, where each isolation class contained approximately the same number of data points and had a 50% overlap with its neighbouring classes (Fig. 3B).

The conditioning plot shows that the strength of the positive relationship between the unrarefied gamma soundscape richness (log_10_) and island size (ha – log_10_) decreases with increasing isolation. This is suggestive of a negative interaction effect between the island area and isolation. As such, we included a model with an interaction term when fitting linear models in the next section.

### 4.5. Fitting linear models

For model fitting, first we constructed a global model using the following equation:

1. *log*_*10*_*(gamma soundscape richness) ~ log*_*10*_*(island area) + isolation + log*_*10*_*(island area)*isolation*

As we know our predictor variables are correlated with each other (Fig. S6), we checked for multicollinearity between the predictors by calculating the Variance Inflation Factor (VIF) for the model without an interaction term (model 3) using the ‘vif’ function from the ‘car’ R-package (Fox and Weisberg 2019 - version 3.1-0). We observed a VIF of 1.44, which is within the acceptable range to retain both predictor variables in the model (Johnston et al. 2018).

<u>Next, we fitted four other candidate models:</u>

2 *log*_*10*_*(gamma soundscape richness) ~ log*_*10*_*(island area)*
3 *log*_*10*_*(gamma soundscape richness) ~ log*_*10*_*(island area) + isolation*
4 *log*_*10*_*(gamma soundscape richness) ~ isolation*
5 *log*_*10*_*(gamma soundscape richness) ~ 1*

### 4.6. Testing model assumptions

For each of these models, we assessed whether the following assumptions were met: (i) a normal distribution of residuals; (ii) homoscedasticity of residuals; (iii) a zero-mean of residuals; and (iv) independence of residual terms. We found that all models had near-zero mean of residuals and independence of residual terms (Table S2). For model 4, the residuals displayed heteroscedacity, as indicated by the Studentized Breusch-Pagan test. Furthermore, for model 1, the Shapiro-Wilkinson test suggests that the residuals deviate from the assumption of normality slightly, however, this is not confirmed by the Kolmogorov-Smirnov test (Table S2; Fig. S7).

**Figure S7:**
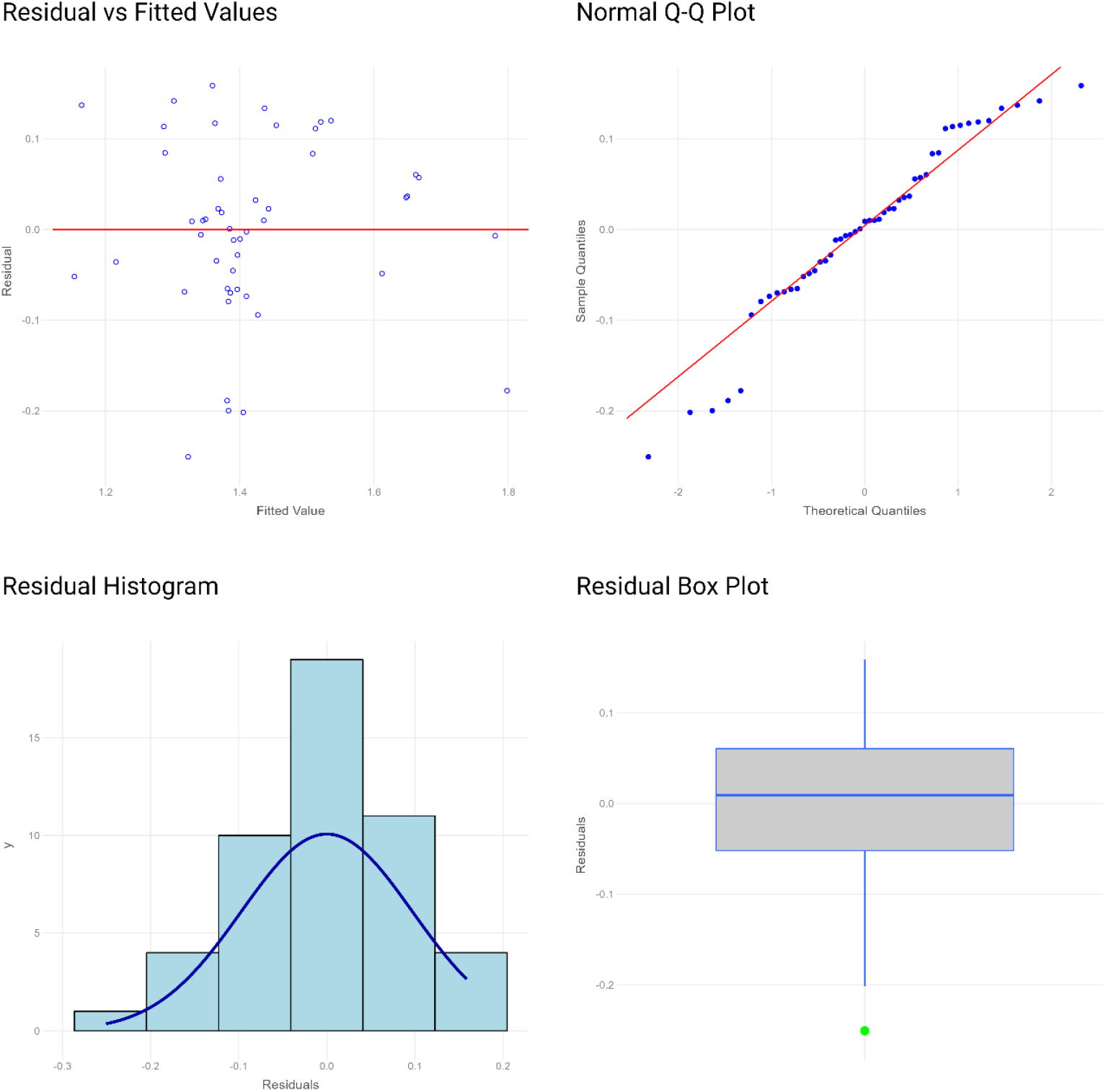
Diagnostic plots showing the residual-vs-fitted plot, residuals qq-plot, residuals boxplot and residuals histogram in a clockwise order from the top left to the bottom left for the best fitting model (model 1).

**Figure S8:**
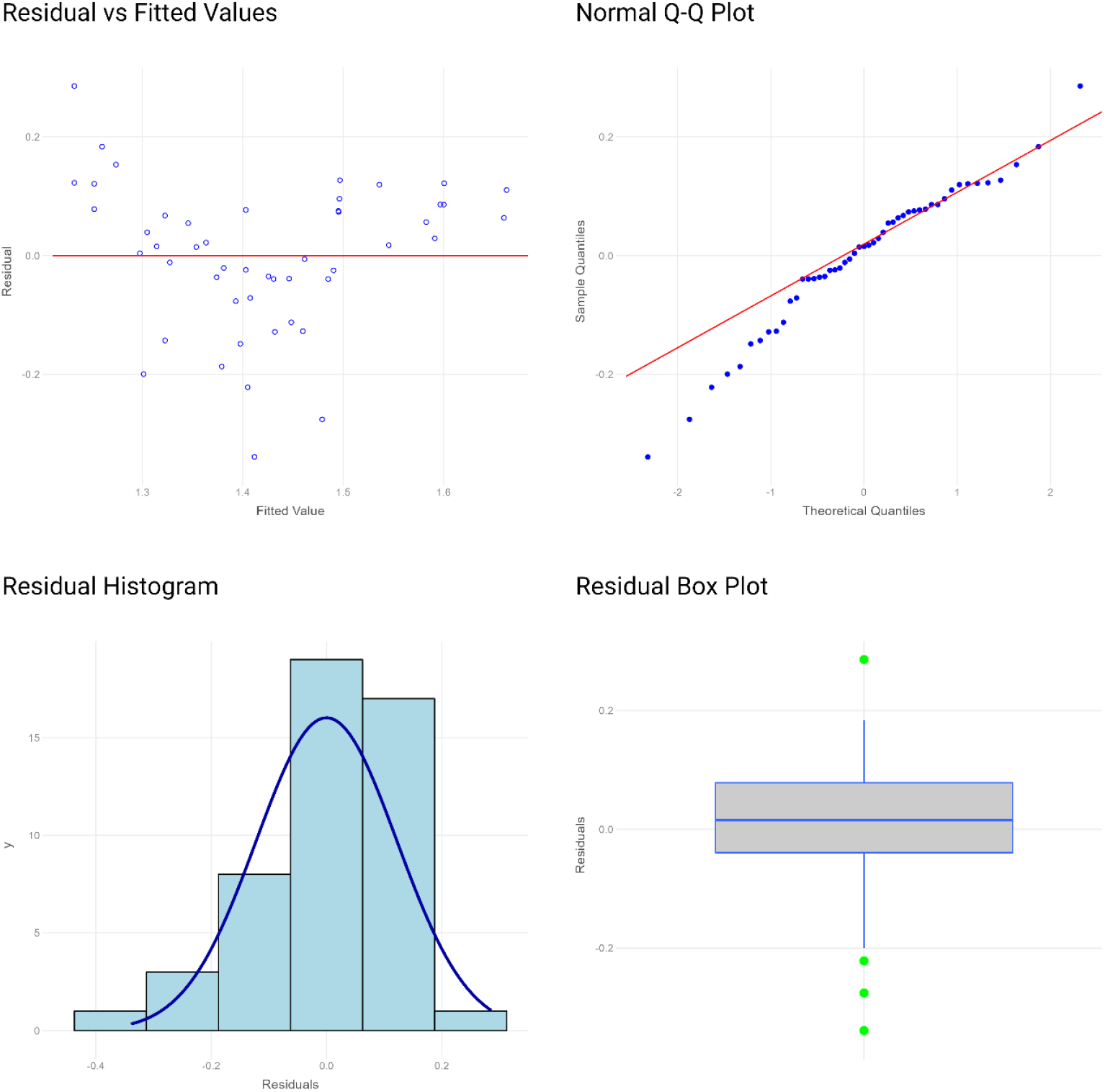
Diagnostic plots showing the residual-vs-fitted plot, residuals qq-plot, residuals boxplot and residuals histogram in a clockwise order from the top left to the bottom left for the second-best fitting model (model 2).

**Table S2:**
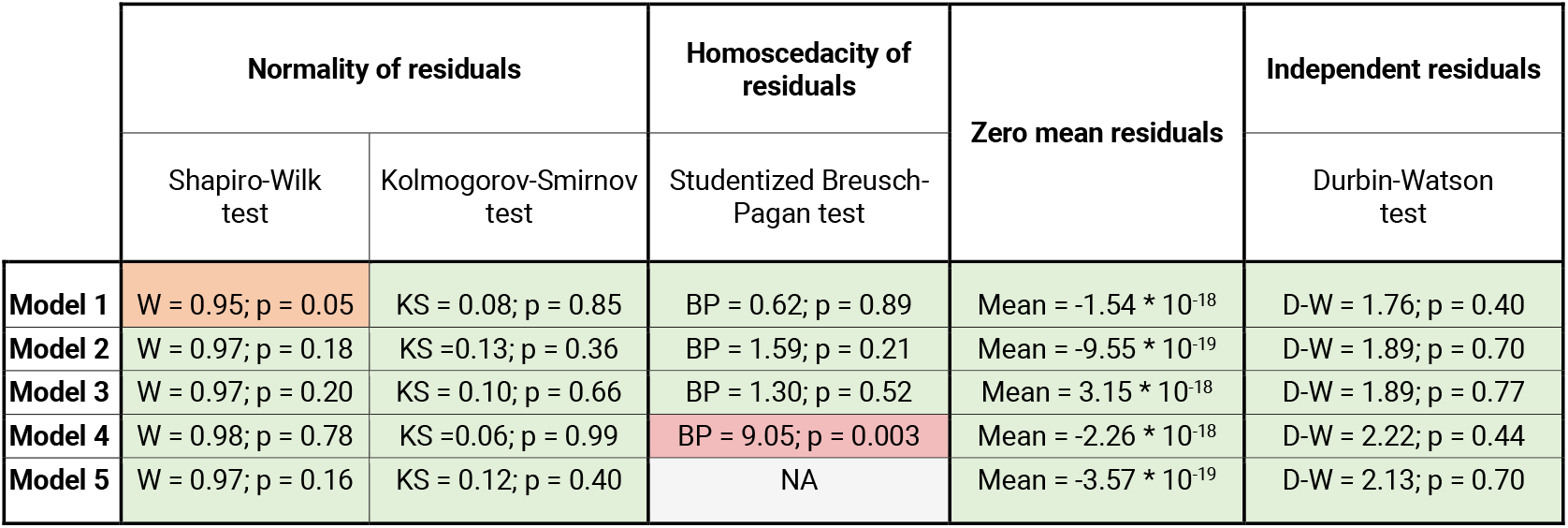
A diagnostic table for five fitted models with results for the following model assumption tests: (i) normal distribution of residuals; (ii) homoscedacity of residuals; (iii)a zero-mean of residuals; and (iv) independence of residuals terms.

## Supplementary material 5: The small-island effect

The perceived slope of the species-area relationship is sensitive to the scale at which ISARs are considered. According to Lomolino and Weiser (2001), untransformed ISARs should exhibit a sigmoidal form with three distinct phases: (i) at very small spatial scales, a phase with no strong relationship between the species richness and island size (the small-island effect or SIE); (ii) at intermediate scales, a phase with a rapid rise in richness with increasing island size; and (iii) at large scales, a phase with a flattening of the slope towards an asymptote, as the number of species per island reaches the levels of the mainland species pool. Each of these phases is delineated by a turnover point at which the predominant mechanisms that govern species richness in space change.

For our assessment of potential soundscape-area relationships and the mechanisms driving them, we were only interested in the effect of the island area on soundscape richness (phase 2). As such, we tested whether there was a spatial scale below which the effect of island area on the soundscape richness broke down (*the small island effect)*. To do so, we used breakpoint linear regression, following the criteria outlined in Dengler (2010) for robust SIE detection: (i) a goodness-of-fit measure that penalized the model complexity; (ii) inclusion of at least three relevant SIE models (a linear, left-horizontal and continuous one-threshold model); (iii) model selection in the same S-space; and (iv) inclusion of islands with zero-richness. We employed the *‘sar_threshold’* function in the ‘sars’ R-package (Matthews et al. 2019) to fit four SIE models: (i) a continuous one-threshold model; (ii) a left-horizontal one-thresold model; (ii) a log-linear model (log_10_); and (iv) an intercept-only model. As the scales considered in this study ranged from small to intermediate, and to avoid overfitting, we did not test for the presence of phase 3 using two-threshold models. We selected the best model fit while penalizing for added model complexity by considering the sample-size-correct Akaike Information Criterion (*AICc*), Bayesian Information Criterion (*BIC*) and the adjusted R^2^ value.

Considering all model selection factors, we found a comparable fit for the continuous one-threshold and left-horizontal one-threshold models with thresholds at 9.40 and 12.68 ha respectively (Table S3; Fig S9). As such, we used the smaller of both threshold values (threshold = 9.40 ha) as our cut-off for the small island effect in our study. Thus, for all subsequent analyses investigating species-area patterns, we will include only the islands above 9.40 ha.

**Table S3:**
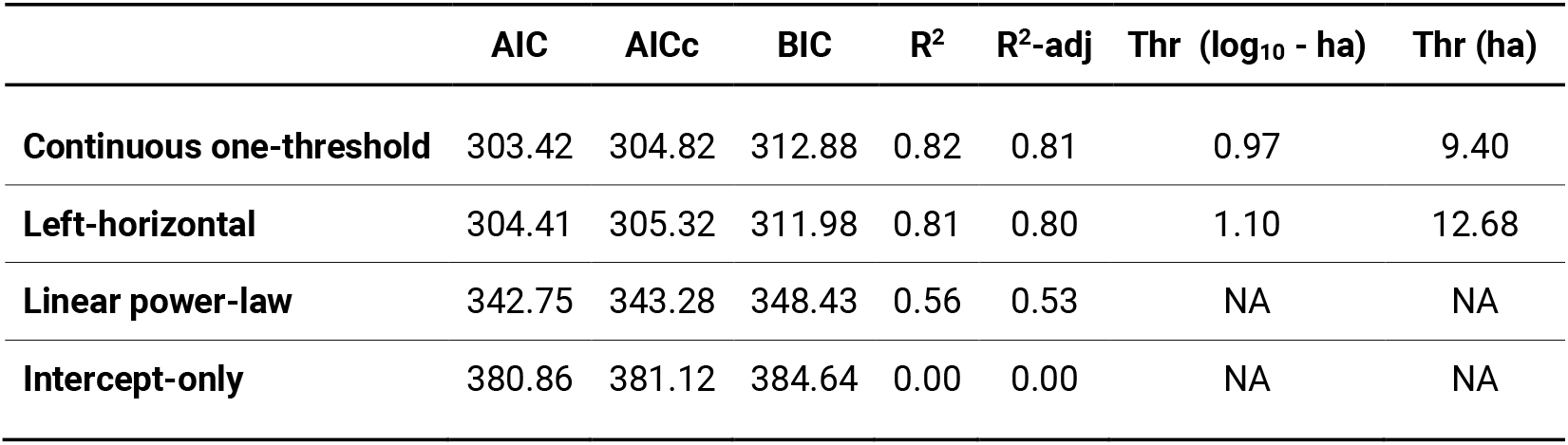
A table containing the SIE model output.

**Figure S9:**
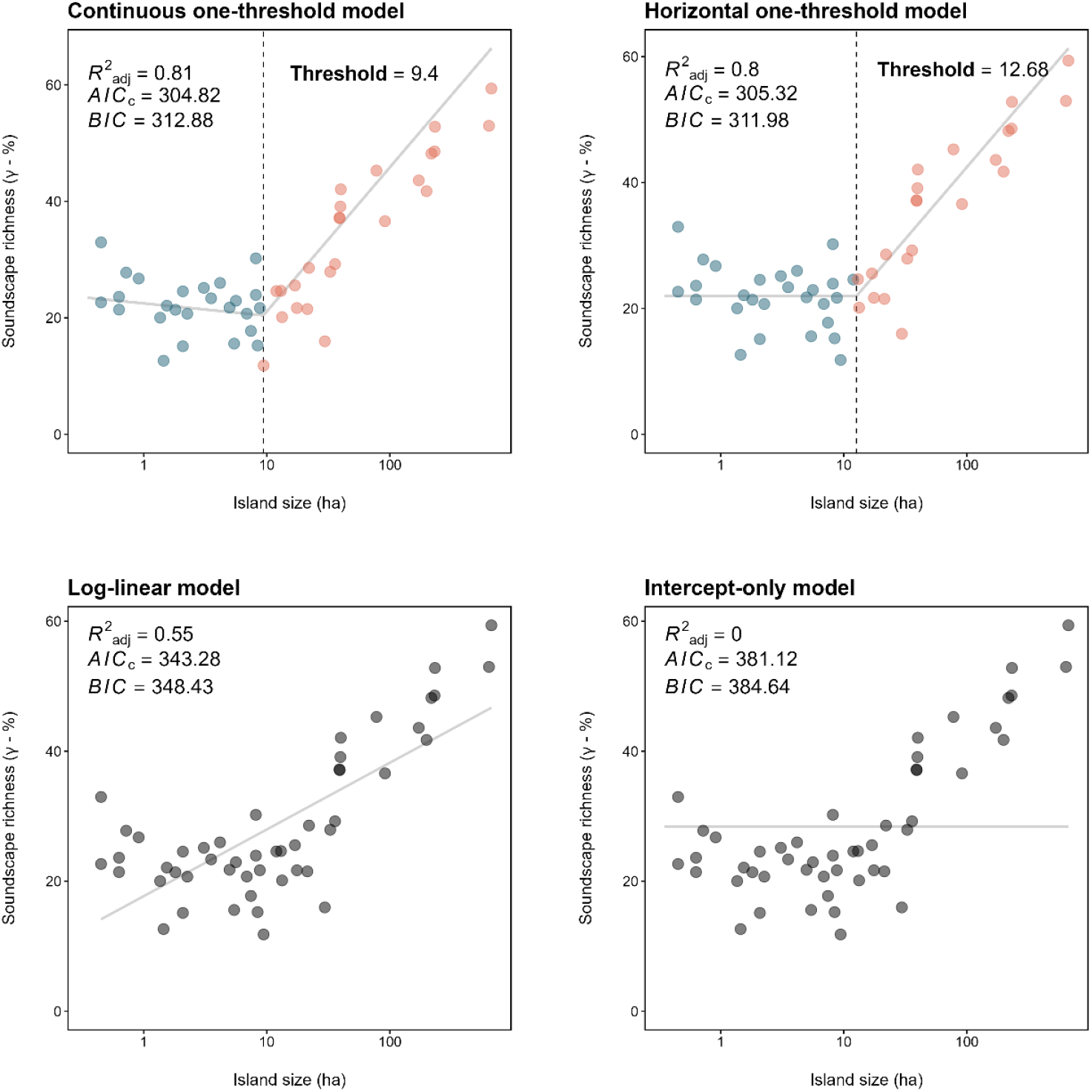
A series of scatterplots with fitted lines (light grey) showing the relationship between the unrarefied gamma soundscape richness and island size (log_10_) for four models: (i) a continuous one-threshold model; (ii) a horizontal one-threshold model; (iii) a log-linear model; and (iv) and intercept-only model. The blue and orange dots indicate data points below and above the model’s threshold respectively, with the dashed black line indicating the island size threshold for that model. Grey dots indicate the model did not include any threshold.

