## supplementary_materials for "Extending species-area relationships into the realm of ecoacoustics: The soundscape-area relationship"

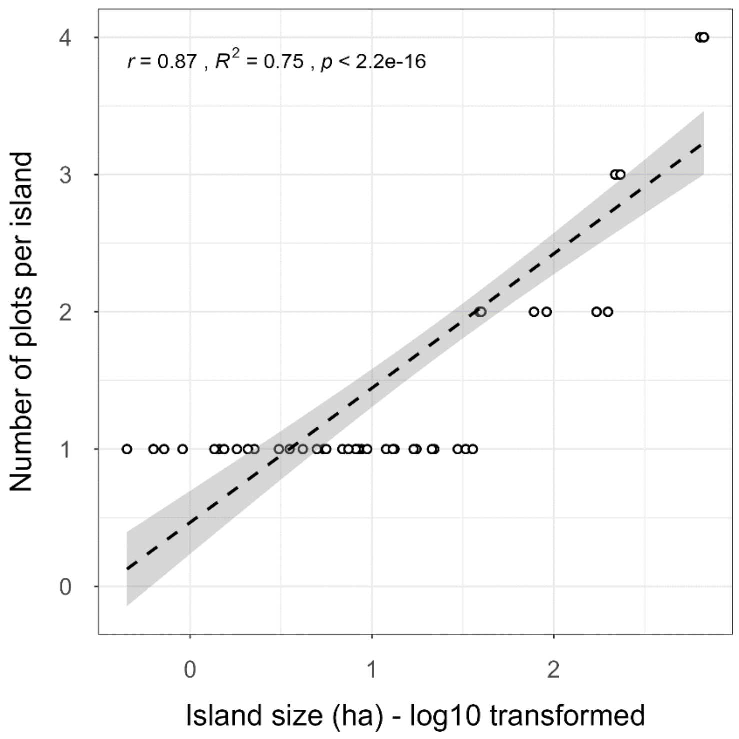


*Figure S1: A scatterplot with regression line showing the proportional relationship between the island size ($log_{10}$ transformed) and the number of acoustic sampling plots per island ($r = 0.87$; $R^2 = 0.75$; $p < 0.005$) for the final set of islands used in the study.*

***Table S1****: An overview of the sites included in the study*

| Island name | Plot name | Latitude | Longitude | Area (ha) |
|---|---|---|---|---|
| 10_626 | 10_626 | -1.75309 | -59.39003 | 8.78 |
| 10_709 | 10_709 | -1.80791 | -59.46624 | 5.43 |
| 12_28 | 12_28 | -1.85177 | -59.4038 | 8.42 |
| 12_9 | 12_9 | -1.5057 | -59.79838 | 11.96 |
| 15_188 | 15_188 | -1.55566 | -59.72653 | 1.81 |
| 17_697 | 17_697 | -1.44519 | -59.8948 | 17.57 |
| 185_358 | 185_358_A | -1.53759 | -59.71342 | 171.73 |
| | 185_358_B | -1.53013 | -59.71838 | 171.73 |
| 2_258 | 2_258 | -1.85392 | -59.38668 | 0.72 |
| 2_333 | 2_333 | -1.83704 | -59.46564 | 0.63 |
| 2_771 | 2_771 | -1.82046 | -59.44777 | 0.45 |
| 2_794 | 2_794 | -1.86325 | -59.39397 | 0.91 |
| 2_87 | 2_87 | -1.7457 | -59.40742 | 1.45 |
| 235_234 | 235_234_A | -1.4518 | -59.81887 | 230.7 |
| | 235_234_B | -1.46164 | -59.81255 | 230.7 |
| | 235_234_C | -1.46213 | -59.82167 | 230.7 |
| 3_311 | 3_311 | -1.87406 | -59.41093 | 1.36 |
| 33_793 | 33_793_A | -1.53047 | -59.7877 | 29.62 |
| 34_526 | 34_526_A | -1.86134 | -59.4132 | 5.61 |
| 37_028 | 37_028_B | -1.77863 | -59.38195 | 32.78 |
| 37_7 | 37_7_A | -1.5103 | -59.79834 | 39.67 |
| | 37_7_B | -1.5063 | -59.80247 | 39.67 |
| 4_746 | 4_746 | -1.54811 | -59.73888 | 2.26 |
| 43_792 | 43_792_A | -1.73547 | -59.47663 | 38.94 |
| | 43_792_B | -1.74108 | -59.47684 | 38.94 |
| 44_174 | 44_174_A | -1.75791 | -59.4764 | 39.12 |
| | 44_174_B | -1.766 | -59.48114 | 39.12 |
| 44_21 | 44_21_B | -1.38776 | -59.76615 | 35.87 |
| 49_62 | 49_62_A | -1.41565 | -59.80996 | 39.94 |
| | 49_62_B | -1.41927 | -59.80871 | 39.94 |
| 5_708 | 5_708 | -1.57007 | -59.71667 | 3.53 |
| 54_544 | 54_544_B | -1.85262 | -59.36253 | 22.01 |
| 7_335 | 7_335 | -1.8555 | -59.4506 | 4.17 |
| 8_042 | 8_042 | -1.54737 | -59.85917 | 6.88 |
| 8_672 | 8_672 | -1.3854 | -59.81981 | 8.15 |

| Island name | Plot name | Latitude | Longitude | Area (ha) |
|---|---|---|---|---|
| Abandonada_left | Abandonada_left | -1.39418 | -59.78186 | 8.15 |
| Abandonada_right | Abandonada_right | -1.39513 | -59.77795 | 0.45 |
| Abusado | Abusado | -1.76217 | -59.67862 | 13.31 |
| Aline | Aline | -1.54838 | -59.75128 | 2.08 |
| Andre | Andre | -1.58462 | -59.87211 | 2.08 |
| Arrepiado | Arrepiado | -1.51483 | -59.73933 | 7.43 |
| Beco_do_Catitu | Beco_do_Catitu_A | -1.75116 | -59.70799 | 638.66 |
| | Beco_do_Catitu_B | -1.73938 | -59.70495 | 638.66 |
| | Beco_do_Catitu_D | -1.73166 | -59.69366 | 638.66 |
| | Beco_do_Catitu_E | -1.72938 | -59.703 | 638.66 |
| Cipoal | Cipoal_A | -1.69814 | -59.78502 | 217.63 |
| | Cipoal_B | -1.70343 | -59.7876 | 217.63 |
| | Cipoal_C | -1.70606 | -59.78051 | 217.63 |
| Coata | Coata | -1.48819 | -59.7872 | 16.94 |
| Formiga | Formiga | -1.83339 | -59.42093 | 1.54 |
| Furo_do_Santa_Luzia | Furo_de_Santa_Luzia_B | -1.74046 | -59.44223 | 198.52 |
| | Furo_de_Santa_Luzia_C | -1.73941 | -59.44632 | 198.52 |
| Garrafa | Garrafa | -1.58863 | -59.83535 | 9.42 |
| Jabuti | Jabuti_A | -1.62741 | -59.76267 | 232.49 |
| | Jabuti_B | -1.62615 | -59.75668 | 232.49 |
| | Jabuti_C | -1.6291 | -59.75201 | 232.49 |
| Joaninha | Joaninha | -1.52243 | -59.82888 | 0.63 |
| Mascote | Mascote_A1 | -1.64506 | -59.82035 | 668.03 |
| | Mascote_A2 | -1.6489 | -59.83297 | 668.03 |
| | Mascote_B1 | -1.64406 | -59.84817 | 668.03 |
| | Mascote_B2 | -1.65936 | -59.83546 | 668.03 |
| Moita | Moita_A | -1.55491 | -59.894 | 91.3 |
| | Moita_B | -1.55799 | -59.89637 | 91.3 |
| Palhal | Palhal | -1.79071 | -59.44785 | 21.37 |
| Panema | Panema | -1.7745 | -59.69236 | 3.08 |
| Pe_Torto | Pe_Torto | -1.76671 | -59.36338 | 4.98 |
| Piquia | Piquia | -1.50662 | -59.78885 | 13.04 |
| Sapupara | Sapupara_A | -1.69722 | -59.61253 | 77.8 |
| | Sapupara_B | -1.69699 | -59.61744 | 77.8 |

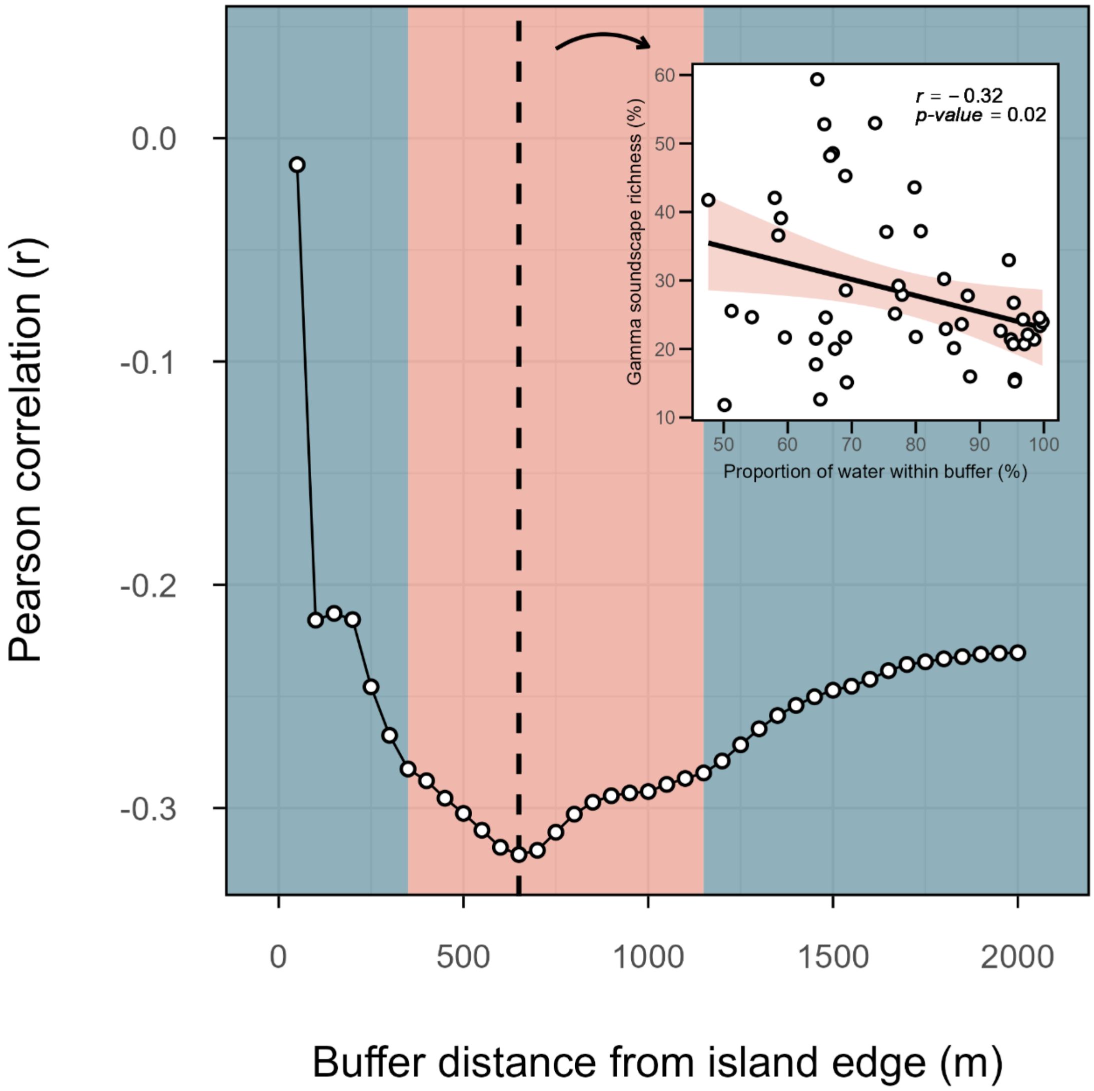


***Figure S2****: A visual representation of the spatial scale at which the landscape-scale isolation metric derived in this study (proportion of water within the buffer area of each island) attains the highest correlation value with the soundscape richness metric (unrarefied gamma soundscape richness); this is better known as the 'scale-of-effect' (Jackson and Fahrig 2015). The blue shading represents buffer distances for which the correlation between soundscape richness and the isolation metric were not significant (p > 0.05), whereas orange shading represents significant correlations. The highest correlation value was reached at a buffer distance of 650 m. The inset plot displays the correlation between the soundscape richness and our isolation metric at the scale-of-effect (650 m).*

The highest correlation between soundscape richness and our isolation metric was attained at a scale-of-effect of 650 m (Fig. S2). At this spatial scale, we observed a significant negative correlation between the degree of island isolation and soundscape richness ($r = -0.32$; $p < 0.05$).

### Supplementary Material 4: Modelling island size versus isolation

To examine the effect of island size versus island isolation on the unrarefied island-wide gamma soundscape richness, we fitted a series of linear models and used an information theoretic approach for model selection (Burnham and Anderson 2004).

##### *4.1. Checking variable distributions*

Before modelling the data, we assessed the distribution of the three variables under investigation (unrarefied gamma soundscape richness, island size, and island isolation) using raincloud plots and Quantile-Quantile plots (Fig. S3, S4 and S5). We observed a right-skewed distribution for the gamma soundscape richness and island area (Fig. S3 and S4). To account for this, these variables were log-transformed ($\log_{10} x$).

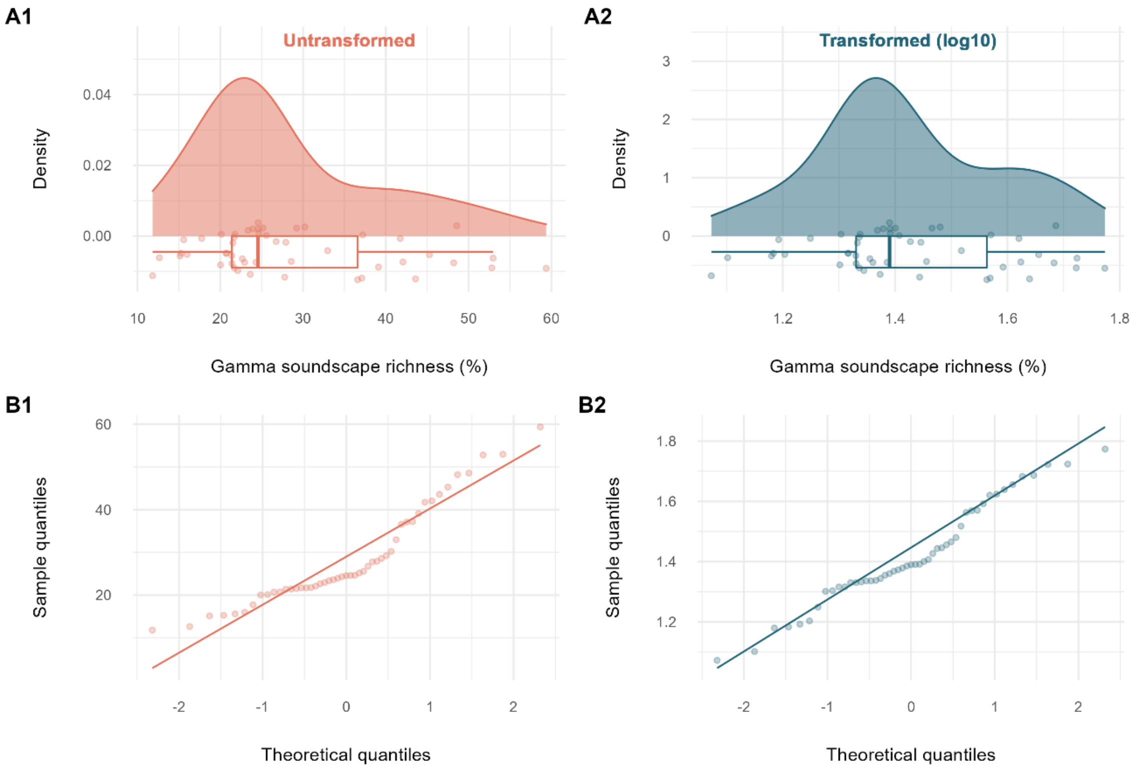


***Figure S3**: A visual representation of the distribution of unrarefied gamma soundscape richness using (A) density plots and (B) quantile-quantile plots for both untransformed (1; orange) and $\log_{10}$-transformed (2; blue) data.*

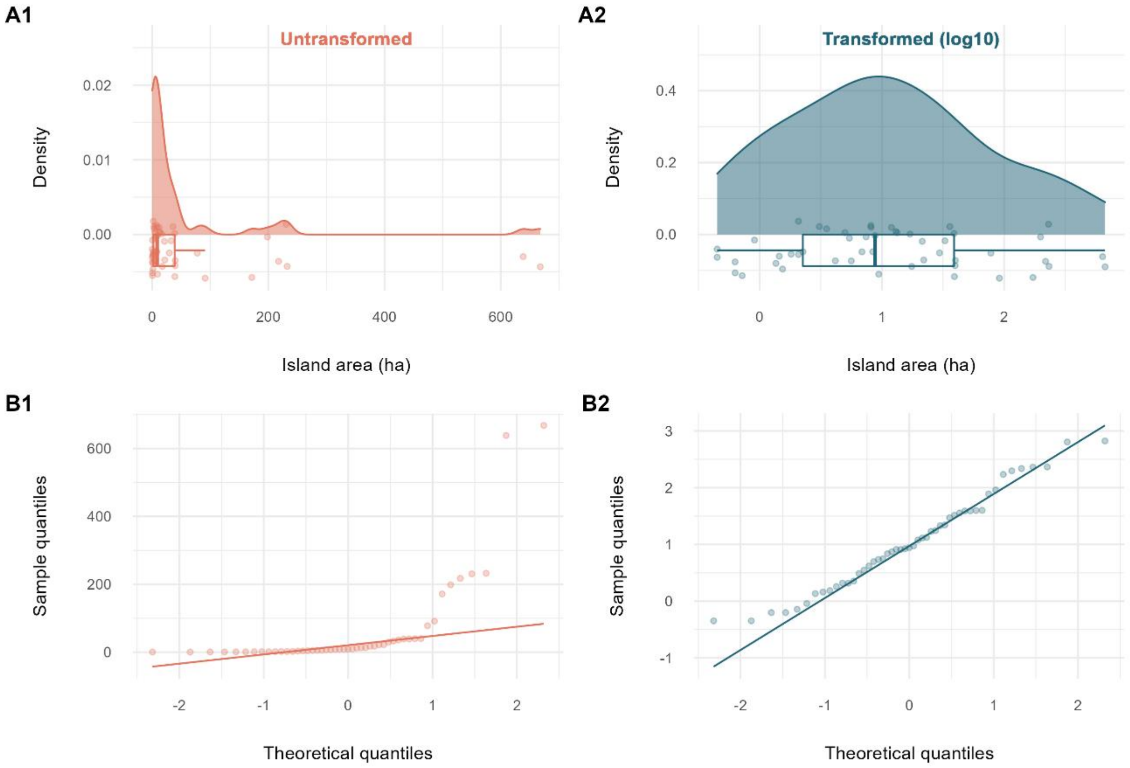


***Figure S4:*** *A visual representation of the distribution of the island size variable using (A) density plots and (B) quantile-quantile plots for both untransformed (1; orange) and* $log_{10}$*-transformed (2; blue) data.*

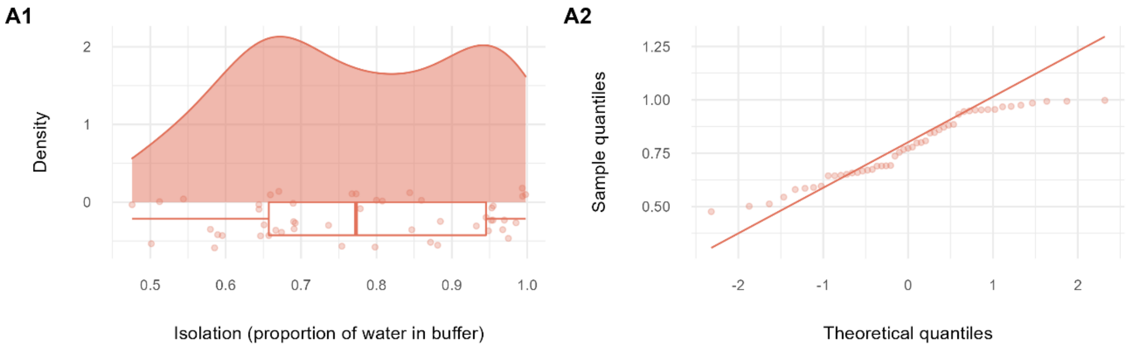


***Figure S5:*** *A visual representation of the distribution of the isolation variable (proportion of water within buffer) using (A1) density plots and (A2) quantile-quantile plots.*

##### *4.2. Exploring the relationship between predictor variables*

In fragmented landscapes, there is often a correlation between the island size and isolation, with smaller islands also being more isolated, or larger islands more connected. As this correlation could have consequences for subsequent modelling procedures, first, we explored the relationship between the size and isolation for the islands contained in the study (Fig. S6).

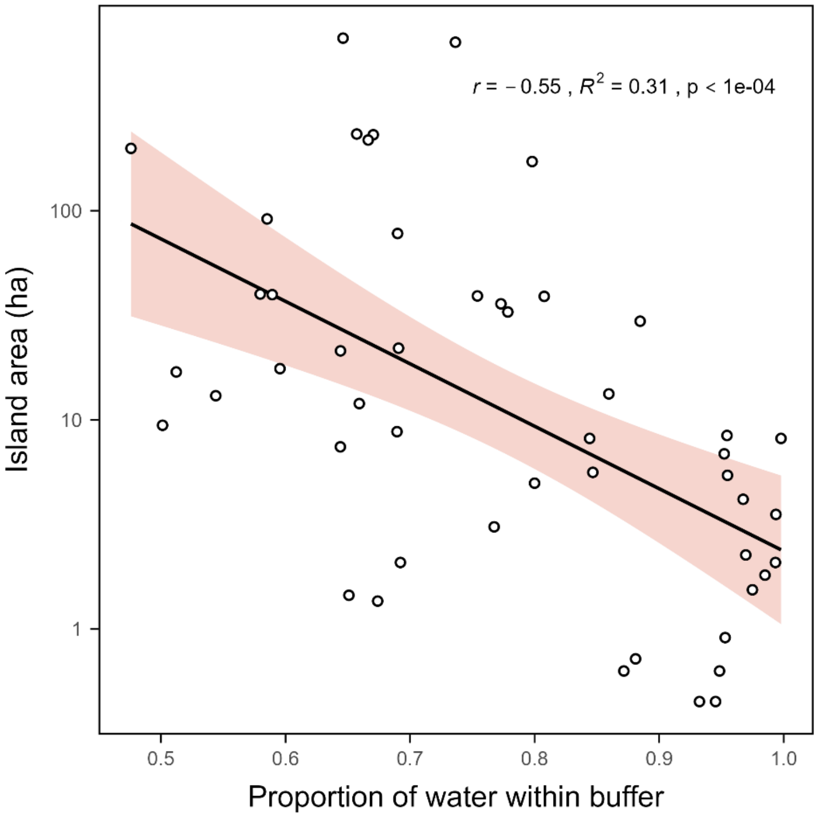


***Figure S6***: *A scatterplot displaying the relationship between the island area (ha – $\log_{10}$ scale) and island isolation (proportion of water within 650 m buffer around the island edge).*

Indeed, as anticipated, our two predictor variables have a strong and negative correlation, a typical observation in fragmented landscapes. Our study system is missing highly connected small islands and highly isolated large islands. We will keep this in the back of our mind as we continue our analysis.

##### *4.5. Fitting linear models*

For model fitting, first we constructed a global model using the following equation:

*(1)* $\log_{10}(\textit{gamma soundscape richness}) \sim \log_{10}(\textit{island area}) + \textit{isolation} + \log_{10}(\textit{island area})*\textit{isolation}$

As we know our predictor variables are correlated with each other (Fig. S6), we checked for multicollinearity between the predictors by calculating the Variance Inflation Factor (VIF) for the model without an interaction term (model 3) using the 'vif' function from the 'car' R-package (Fox and Weisberg 2019 - version 3.1-0). We observed a VIF of 1.44, which is within the acceptable range to retain both predictor variables in the model (Johnston et al. 2018).

Next, we fitted four other candidate models:

*(2)* $\log_{10}(\textit{gamma soundscape richness}) \sim \log_{10}(\textit{island area})$

*(3)* $\log_{10}(\textit{gamma soundscape richness}) \sim \log_{10}(\textit{island area}) + \textit{isolation}$

*(4)* $\log_{10}(\textit{gamma soundscape richness}) \sim \textit{isolation}$

*(5)* $\log_{10}(\textit{gamma soundscape richness}) \sim 1$

##### *4.6. Testing model assumptions*

For each of these models, we assessed whether the following assumptions were met: (i) a normal distribution of residuals; (ii) homoscedasticity of residuals; (iii) a zero-mean of residuals; and (iv) independence of residual terms. We found that all models had near-zero mean of residuals and independence of residual terms (Table S2). For model 4, the residuals displayed heteroscedacity, as indicated by the Studentized Breusch-Pagan test. Furthermore, for model 1, the Shapiro-Wilkinson test suggests that the residuals deviate from the assumption of normality slightly, however, this is not confirmed by the Kolmogorov-Smirnov test (Table S2; Fig. S7).

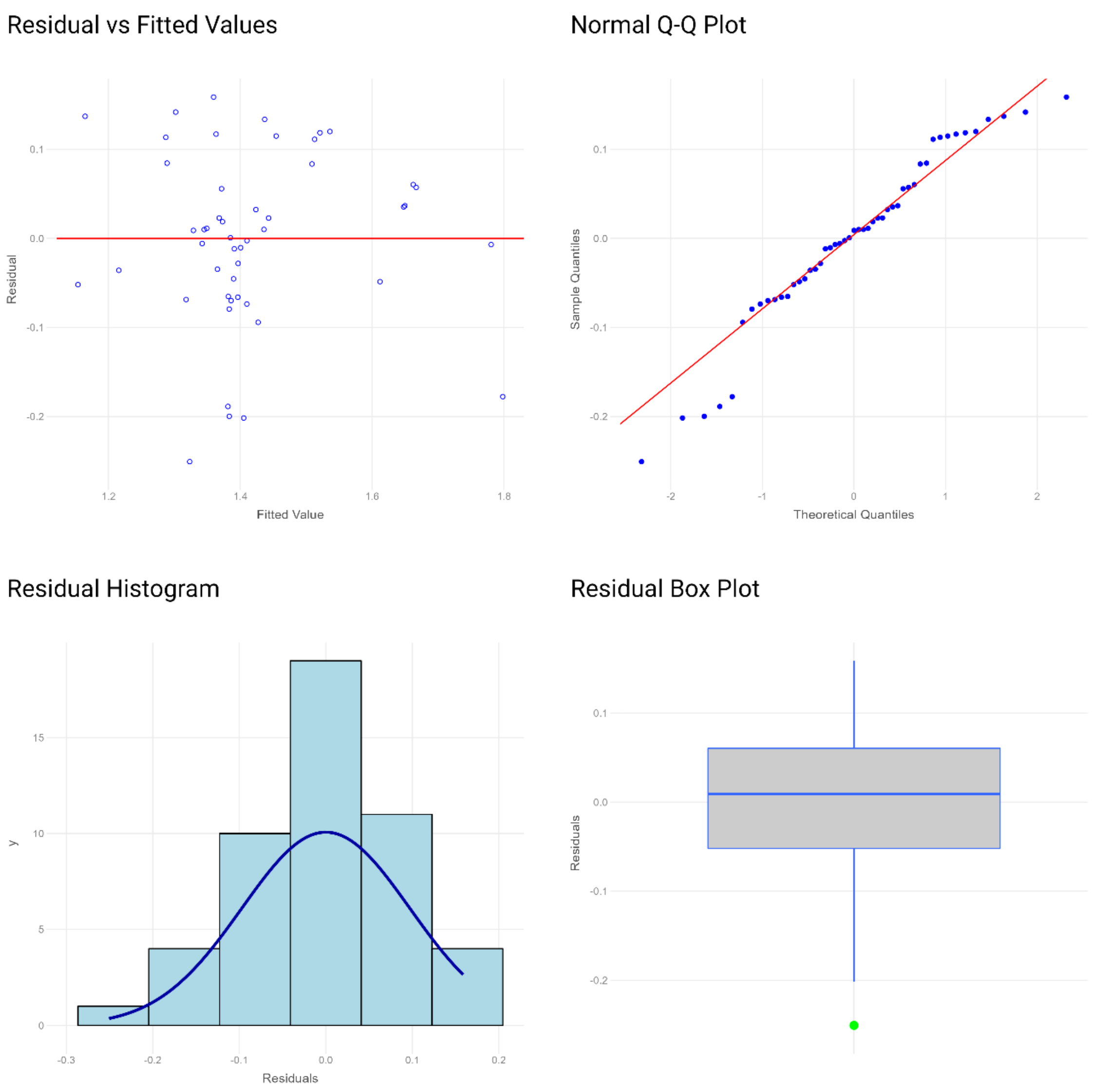


***Figure S7***: *Diagnostic plots showing the residual-vs-fitted plot, residuals qq-plot, residuals boxplot and residuals histogram in a clockwise order from the top left to the bottom left for the best fitting model (model 1).*

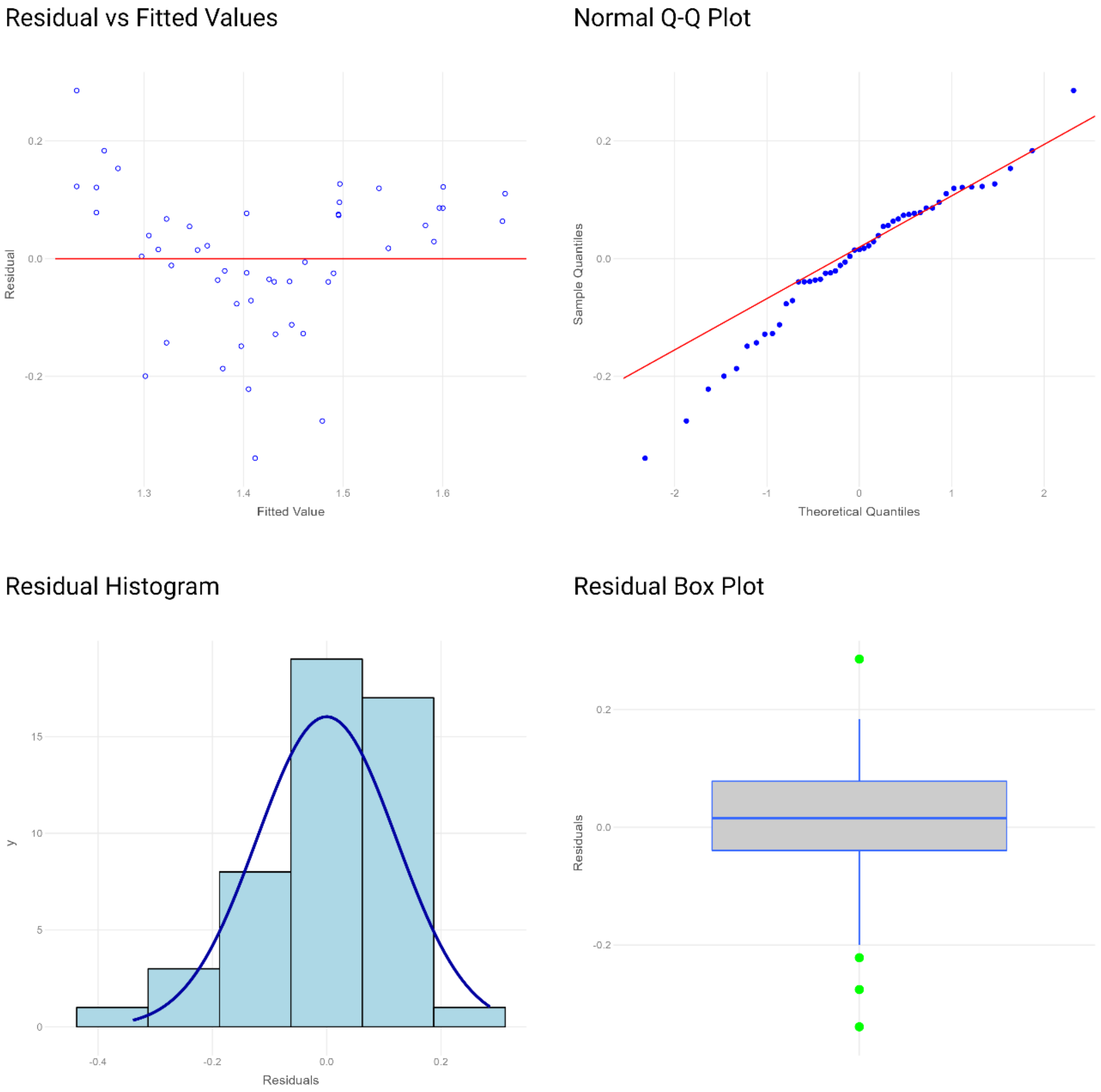


***Figure S8***: *Diagnostic plots showing the residual-vs-fitted plot, residuals qq-plot, residuals boxplot and residuals histogram in a clockwise order from the top left to the bottom left for the second-best fitting model (model 2).*

***Table S2**: A diagnostic table for five fitted models with results for the following model assumption tests: (i) normal distribution of residuals; (ii) homoscedacity of residuals; (iii)a zero-mean of residuals; and (iv) independence of residuals terms.*

| | **Normality of residuals** | | **Homoscedacity of residuals** | **Zero mean residuals** | **Independent residuals** |
|---|---|---|---|---|---|
| | Shapiro-Wilk test | Kolmogorov-Smirnov test | Studentized Breusch-Pagan test | | Durbin-Watson test |
| **Model 1** | W = 0.95; p = 0.05 | KS = 0.08; p = 0.85 | BP = 0.62; p = 0.89 | Mean = -1.54 * $10^{-18}$ | D-W = 1.76; p = 0.40 |
| **Model 2** | W = 0.97; p = 0.18 | KS =0.13; p = 0.36 | BP = 1.59; p = 0.21 | Mean = -9.55 * $10^{-19}$ | D-W = 1.89; p = 0.70 |
| **Model 3** | W = 0.97; p = 0.20 | KS = 0.10; p = 0.66 | BP = 1.30; p = 0.52 | Mean = 3.15 * $10^{-18}$ | D-W = 1.89; p = 0.77 |
| **Model 4** | W = 0.98; p = 0.78 | KS =0.06; p = 0.99 | BP = 9.05; p = 0.003 | Mean = -2.26 * $10^{-18}$ | D-W = 2.22; p = 0.44 |
| **Model 5** | W = 0.97; p = 0.16 | KS = 0.12; p = 0.40 | NA | Mean = -3.57 * $10^{-19}$ | D-W = 2.13; p = 0.70 |

***Table S3**: A table containing the SIE model output.*

| | AIC | AICc | BIC | $R^2$ | $R^2$-adj | Thr (log$_{10}$ - ha) | Thr (ha) |
|---|---|---|---|---|---|---|---|
| **Continuous one-threshold** | 303.42 | 304.82 | 312.88 | 0.82 | 0.81 | 0.97 | 9.40 |
| **Left-horizontal** | 304.41 | 305.32 | 311.98 | 0.81 | 0.80 | 1.10 | 12.68 |
| **Linear power-law** | 342.75 | 343.28 | 348.43 | 0.56 | 0.53 | NA | NA |
| **Intercept-only** | 380.86 | 381.12 | 384.64 | 0.00 | 0.00 | NA | NA |

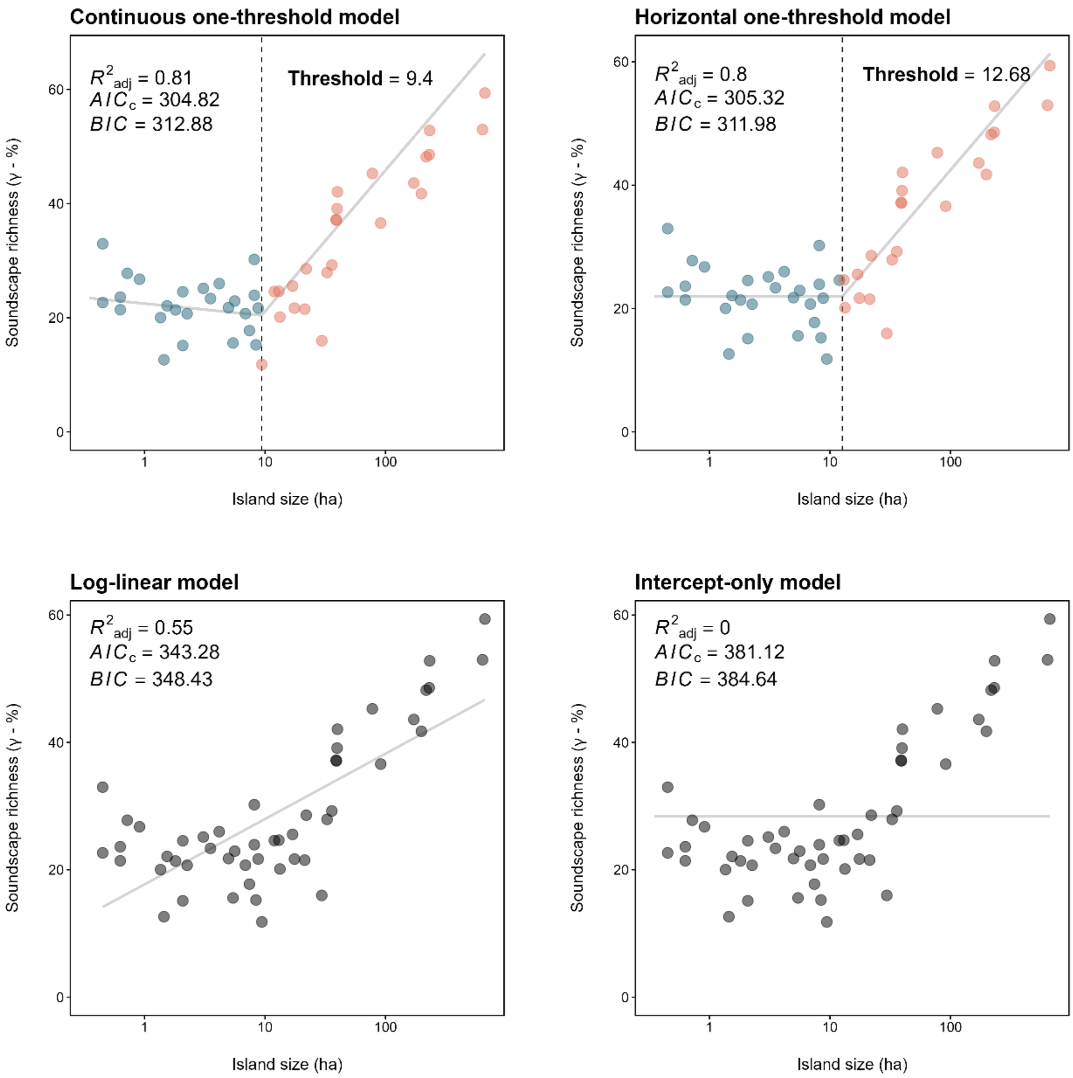


***Figure S9****: A series of scatterplots with fitted lines (light grey) showing the relationship between the unrarefied gamma soundscape richness and island size ($log_{10}$) for four models: (i) a continuous one-threshold model; (ii) a horizontal one-threshold model; (iii) a log-linear model; and (iv) and intercept-only model. The blue and orange dots indicate data points below and above the model's threshold respectively, with the dashed black line indicating the island size threshold for that model. Grey dots indicate the model did not include any threshold.*
